# Genomic novelty and process-level convergence in adaptation to whole genome duplication

**DOI:** 10.1101/2020.01.31.929109

**Authors:** Magdalena Bohutínská, Mark Alston, Patrick Monnahan, Terezie Mandáková, Sian Bray, Pirita Paajanen, Filip Kolář, Levi Yant

## Abstract

Whole genome duplication (WGD) occurs across kingdoms and can promote adaptation. However, a sudden increase in chromosome number, as well as changes in physiology, are traumatic to conserved processes. Previous work in *Arabidopsis arenosa* revealed a coordinated genomic response to WGD, involving physically interacting meiosis proteins, as well as changes related to cell cycle and ion homeostasis. Here we ask: is this coordinated shift in the same processes repeated in another species following WGD? To answer this, we resequenced and cytologically assessed replicated populations from a diploid/autotetraploid system, *Cardamine amara*, and test the hypothesis that gene and process-level convergence will be prevalent between these two WGDs adaptation events. Interestingly, we find that gene-level convergence is negligible, with no more in common than would be expected by chance. This was most clear at meiosis-related genes, consistent with our cytological assessment of somewhat lower meiotic stability in *C. amara*, despite establishment and broad occurrence of the autotetraploid in nature. In contrast, obvious convergence at the level of functional processes, including meiotic cell cycle, chromosome organisation and stress signalling was evident. This indicates that the two autotetraploids survived challenges attendant to WGD via contrasting solutions, modifying different players from similar processes. Overall, this work gives the first insight into the salient adaptations required to cope with a genome-doubled state and brings the first genomic evidence that autopolyploids can utilize multiple trajectories to achieve adaptation to WGD. We speculate that this flexibility increases the likelihood a nascent polyploid overcomes early stringent challenges to later access the spectrum of evolutionary opportunities of polyploidy.

**Significance statement:** Whole genome duplication (WGD) is a tremendous mutation and an important evolutionary force. It also presents immediate changes to meiosis and cell physiology that nascent polyploids must overcome to survive. Given the dual facts that WGD adaptation is difficult, but many lineages nevertheless survive WGD, we ask: how constrained are the evolutionary responses to a genome-doubled state? We previously identified candidate genes for WGD adaptation in *Arabidopsis arenosa*, which has natural diploid and tetraploid variants. Here we test for evolutionary convergence in adaptation to WGD in a species 17 million years distant, *Cardamine amara*. This work gives the first genomic insight into of how autopolyploids utilize multiple adaptive trajectories to manage a genome-doubled state.

## Introduction

Whole genome duplication (WGD) is both a massive mutation and a powerful force in evolution (1). The opportunities and challenges presented by WGD occur immediately, realised in a single generation. As such, WGD comes as a shock to the system. Autopolyploids, formed by within-species WGD (without hybridization), emerge from the chance encounter of unreduced gametes. Thus, they typically harbour four full haploid genomes that are similar in all pairwise combinations, resulting in a lack of pairing partner preferences at meiosis. This, combined with multiple crossover events per chromosome pair, can result in entanglements among three or more homologs at anaphase and mis-segregation or chromosome breakage, leading to aneuploidy (2–4). Beyond this, WGD presents a suddenly transformed intracellular landscape to the conserved workings of the cell, such as altered ion homeostasis and a host of nucleotypic factors related to cell size, volume, and cell cycle progression (3, 5, 6). Occasionally however, a lineage survives this early trauma and graduates to runaway evolutionary success. Indeed, there is some direct empirical evidence of the increased adaptability of autopolyploid lineages from *in vitro* evolutionary competition experiments (7). With increased ploidy, genetic variability can be maintained in a masked state, with evidence of lineages acting as allelic sponges recruiting diverse alleles by gene flow across ploidies, and indeed, species (8, 9). Thus, while substantial opportunities await lineages that adapt to WGD, clear challenges must be overcome to function as a polyploid (3, 10, 11).

The genomic basis for adaptation to WGD has been most extensively investigated in *Arabidopsis arenosa*, which exists as both diploid and autotetraploid in the wild (12). There, the strongest genomic signals of adaptation to WGD are in a suite of 8 genes that cooperatively govern early events in the formation of meiotic chromosome crossovers (13, 14). The products of these genes physically and functionally interact to control this coordinated, conserved process, which stands as a leading candidate process mediating adaptation to WGD. In the evolved *A. arenosa* autotetraploids harbouring these selected alleles, we observed a decrease in meiotic crossover number as well as fewer chromosome entanglements relative to synthetic autopolyploids with ancestral diploid alleles (14). Recent work found that the sister species *Arabidopsis lyrata*, a younger autotetraploid, also harbours many of the same selected alleles discovered in *A. arenosa* (9). Moreover, from a joint population genomic analysis of both species across a hybrid zone, clear signals of bidirectional adaptive gene flow emerge exactly at these adaptive alleles between *A. arenosa* and *A. lyrata* (9, 15). Therefore, *A. lyrata* and *A. arenosa* WGD stabilisation events are not independent.

Here we use an independent system, ~17 million years diverged from both *A. arenosa* and *A. lyrata* (16), to test the hypothesis that this solution of nimble meiosis gene evolution is repeated, and if not, whether changes in other genes from analogous processes are associated with adaptation to WGD. Given the clear results in *A. arenosa* and *A. lyrata*, we hypothesised that the adaptive trajectories which are available to mediate adaptation to a WGD state may be constrained, leading to repeated selection of the same meiosis genes. Such a result would offer a striking case of convergent evolution in core cellular processes. To test this hypothesis, we take advantage of a well-characterised model, *Cardamine amara* (Brassicaceae, tribe Cardamineae). A large-scale cytotyping survey and genetic analysis demonstrated an autopolyploid origin of the successful autotetraploid cytotype found in the Eastern and Central Alps (17–20). Importantly, *C. amara* is a perennial herb harbouring a high level of genetic diversity (similar to both *A. arenosa* and *A. lyrata*) and shares a similar distribution range and evolutionary history, with a likely single origin, followed by autotetraploid expansion associated with glacial oscillations (18, 19).

To test our hypothesis of gene and process-level convergence, we performed genome scans for selection, contrasting natural autotetraploid and diploid populations. We found the strongest selection signals specifically at genes involved in cellular functions central to adaptation to WGD: chromosome remodelling, meiosis, cell cycle regulation, and ion transport. However, the evolutionary response to WGD in *C. amara* is very different to that of *A. arenosa*: overall, we saw minimal gene-level convergence in loci putatively mediating adaptation to WGD. In particular, none of the same meiosis-related genes that control meiotic chromosome crossovers in *A. arenosa* were under selection in *C. amara*. This is consistent with observations of clonal spreading and lower meiotic stability in both diploid and autotetraploid *C. amara*, suggesting that *C. amara* autotetraploids are already prepared by their diploid lifestyle to thrive, at least temporarily, while suffering reduced meiotic fidelity. However, in contrast to a lack of gene convergence, we find a strong signal of process-level convergence in core processes controlling DNA management, chromosome organisation, stress signalling, and ion homeostasis. Overall, our results provide sharp contrast to widespread reports of gene-level convergence across the tree of life and suggest that the genomic changes associated with a WGD state might not be as constrained as would be expected based on their functional conservation across eukaryotes.

## Results and Discussion

### Population selection, sampling and genetic structure

To assess the genetic basis of adaptation to WGD in *C. amara*, we generated a novel synthetic long-read reference genome (N50 = 1.82 mb, 95% complete BUSCOs; see Methods) and resequenced in triplicate four populations of contrasting ploidy, sampling 100 individuals: two diploid (LUZ, VRK) and two autotetraploid (CEZ, PIC; Fig. 1A; *SI Appendix*, Table S1). We chose these populations based on a comprehensive cytological survey of over 3,000 *C. amara* samples throughout the Czech Republic (18). The populations we sampled represent core areas of each cytotype, away from potential hybrid zones and distant from any triploid-containing populations. Further, we performed flow cytometry on all samples sequenced to verify expected ploidy.

**Figure 1.**
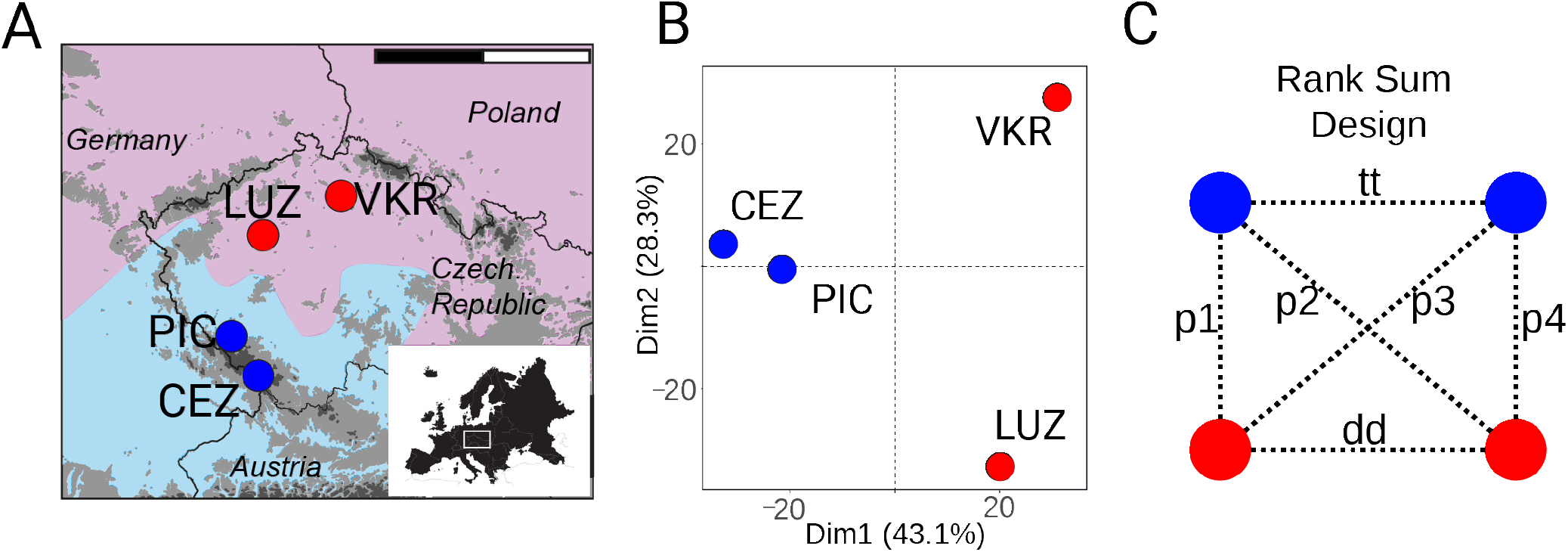
Sample population locations and population structure of *Cardamine amara*. **A**, Locations of *C. amara* populations sampled in the Czech Republic (red, diploids; blue, autotetraploids; scale bar corresponds to 200 km; shaded area represents each cytotype range from (18), with autotetraploid range expansion southward). **B**, Population structure represented by Principal Component Analysis of ~124,000 fourfold degenerate SNPs. **C**, Rank Sum design used to minimise any influence of population-specific divergence in tests for directional selection. ‘p1’ to ‘p4’ represent the between-ploidy contrasts used for the rank sum calculations. ‘dd’ and ‘tt’ represent within-ploidy contrasts used to subtract signal of local population history.

To obtain robust population allele frequency estimates across genomes, we performed a replicated pooled sequencing approach. From each population we pooled DNA from 25 individuals in triplicate and generated on average 31 million reads per pooled sample. We mapped the reads onto our *C. amara* assembly. After read mapping, variant calling and quality filtration, we obtained a final dataset of 2,477,517 SNPs (mean coverage depth per population = 86, *SI Appendix*, Table S2).

Population structure of *C. amara* (Fig. 1B) showed primary differentiation by ploidy (first axis explained 43% of all variability) while the second axis (28% of variability explained) differentiated the two diploid populations from each other. The two autotetraploid populations had the lowest genetic differentiation of all contrasts (Fst = 0.04, mean allele frequency difference = 0.06) and showed a complete absence of fixed differences (Table 1). Close genetic similarity together with spatial arrangement (the populations represent part of a continuous range of autotetraploid cytotype spanning to Eastern Alps) suggest that both autotetraploid populations represent the outcome of a single polyploidization event, in line with previous population genetic inference based on large-scale sampling (18). The similar level of interploidy divergence within both *C. amara* and *A. arenosa* (average Fst between diploids and autotetraploids = 0.10 and 0.11, respectively) suggests that the polyploidization events in both species happened at roughly comparable time points in the past (Table 1).

**Table 1.** Measures of genome-wide differentiation between *C. amara* and *A. arenosa* populations

| Populations | Ploidies | AFD | Fixed diff | Fst | # SNPs |
| --- | --- | --- | --- | --- | --- |
| PIC - VKR | 4x - 2x | 0.09 | 30 | 0.09 | 2,326,315 |
| PIC - LUZ | 4x - 2x | 0.09 | 2 | 0.08 | 2,314,229 |
| CEZ - VKR | 4x - 2x | 0.11 | 120 | 0.12 | 2,333,538 |
| CEZ - LUZ | 4x - 2x | 0.11 | 86 | 0.11 | 2,335,004 |
| CEZ - PIC | 4x - 4x | 0.06 | 0 | 0.04 | 2,297,229 |
| LUZ - VKR | 2x - 2x | 0.1 | 6 | 0.09 | 2,018,892 |
| *A. arenosa* tetraploids - *A. arenosa* diploids | 4x - 2x | 0.05 | 21 | 0.11 | 7,106,848 |

**Note:** Differentiation metrics shown are mean allele frequency difference (AFD), the number of fixed differences (Fixed diff) and $F_{st}$ . In the case of *A. arenosa*, $F_{st}$ in diploids is calculated as a mean over all pairwise $F_{st}$ measurements between the five previously characterised diploid lineages (8).

### Directional selection specifically associated with WGD in *C. amara*

To minimise false positives due to local population history we leveraged a quartet-based sampling design (21), consisting of two diploid and two autotetraploid populations (Fig. 1C). We calculated Fst for 1 kb windows with a minimum 20 SNPs for all six possible population contrasts, and ranked windows based on Fst values. The mean number of SNPs per population contrast was 2,270,868 (Table 1). To focus on WGD-associated adaptation, we first assigned ranks to each window based on the Fst values in each of four possible pairwise diploid-autotetraploid contrasts and identified windows in the top 1% outliers of the resultant combined rank sum (Fig. 1C, contrasts p1-p4). We then excluded any window which was also present in the top 1% Fst outliers in diploid-diploid or autotetraploid-autotetraploid population contrasts to avoid misattribution caused by local population history (Fig. 1C, contrasts ‘tt’ and ‘dd’). By this conservative approach, we identified 440 windows that intersected 229 gene coding loci (*SI Appendix*, Table S3; ‘WGD adaptation candidates’ below). Among these 229 gene coding loci, a Gene Ontology (GO) term enrichment analysis yielded 22 significantly enriched biological processes (Fisher’s exact test with conservative ‘elim’ method, adjusted p < 0.05, *SI Appendix*, Table S4). To further refine the gene list to putatively functional candidates we complemented these differentiation measures with a quantitative estimate following the fineMAV method (22) (see Methods). SNPs were assigned a fineMAV score based on the predicted functional consequences of amino acid substitutions, using Grantham scores, amplified by the allele frequency difference between the two amino acids (22). From our 229 Fst WGD adaptation candidates, 120 contained at least one 1% fineMAV outlier amino acid substitution (*SI Appendix*, Table S3, S5).

### DNA maintenance (repair, chromosome organisation) and meiosis under selection in *C. amara*

Of the 22 significantly enriched GO processes, the most significantly enriched by far was ‘DNA metabolic process’ (p-value = 6.50E-08, vs 0.00021 for the next most confident enrichment), although there was also enrichment for ‘chromosome organization’ and ‘meiotic cell cycle.’ The 40 genes contributing to these categories have specifically localised peaks of differentiation (Fig. 2), as well as 1% fineMAV outlier SNPs in their gene coding regions (Fig. 2, *SI Appendix*, Table S3). These genes also cluster in STRING interaction networks, suggesting coevolutionary dynamics driving these selection signatures (Figure S1; see Methods and Results below). The largest cluster comprises of *MSH6, PDS5e, SMC2, MS5, PKL, HDA18, CRC*, and homologs of two uncharacterised, but putative DNA repair related loci *AT1G52950* and *AT3G02820* (containing SWI3 domain). *MutS Homolog 6* (*MSH6*) is a component of the post-replicative DNA mismatch repair system. It forms a heterodimer with MSH2 which binds to DNA mismatches (23, 24), enhancing mismatch recognition. *MutS* homologs have also been shown to control crossover number in *A. thaliana* (25). The *C. amara* ortholog of *AT1G15940* is a close homolog of *PDS5*, a protein required in fungi and animals for formation of the synaptonemal complex and sister chromatid cohesion (26)*. Structural Maintenance Of Chromosomes 2* (*SMC2/TTN3*) is a central component of the condensin complex, which is required for segregation of homologous chromosomes at meiosis (27) and stable mitosis (28). *PICKLE* (*PKL*) is a SWI/SWF nuclear-localized chromatin remodelling factor (29, 30) that also has highly pleiotropic roles in osmotic stress response (31), stomatal aperture (32), root meristem activity (33), and flowering time (34). Beyond this cluster, other related DNA metabolism genes among our top outliers include *DAYSLEEPER* (Fig. 2), a domesticated transposase that is essential for plant development, first isolated as binding the *Kubox1* motif upstream of the DNA repair gene *Ku70* (35). The complex Ku70/Ku80 regulate non-homologous end joining (NHEJ) double-strand break repair (36). Consistent with this, *DAYSLEEPER* mutants accumulate DNA damage (37), but the exact role of *DAYSLEEPER* in normal DNA maintenance is not understood. Interesting also is the identification of *MALE-STERILE 5* (*MS5/TDM1*), which is required for cell cycle exit after meiosis II. As the name implies, MS5 mutants are male sterile, with pollen tetrads undergoing an extra round of division after meiosis II without chromosome replication (38). *MS5/TDM1* may be an APC/C component whose function is to ensure meiosis termination at the end of meiosis II (39). Together, this set of DNA management loci exhibiting the strongest signals of selection points to widespread modulation of DNA repair and chromosome management following WGD in *C. amara*.

**Figure 2.**
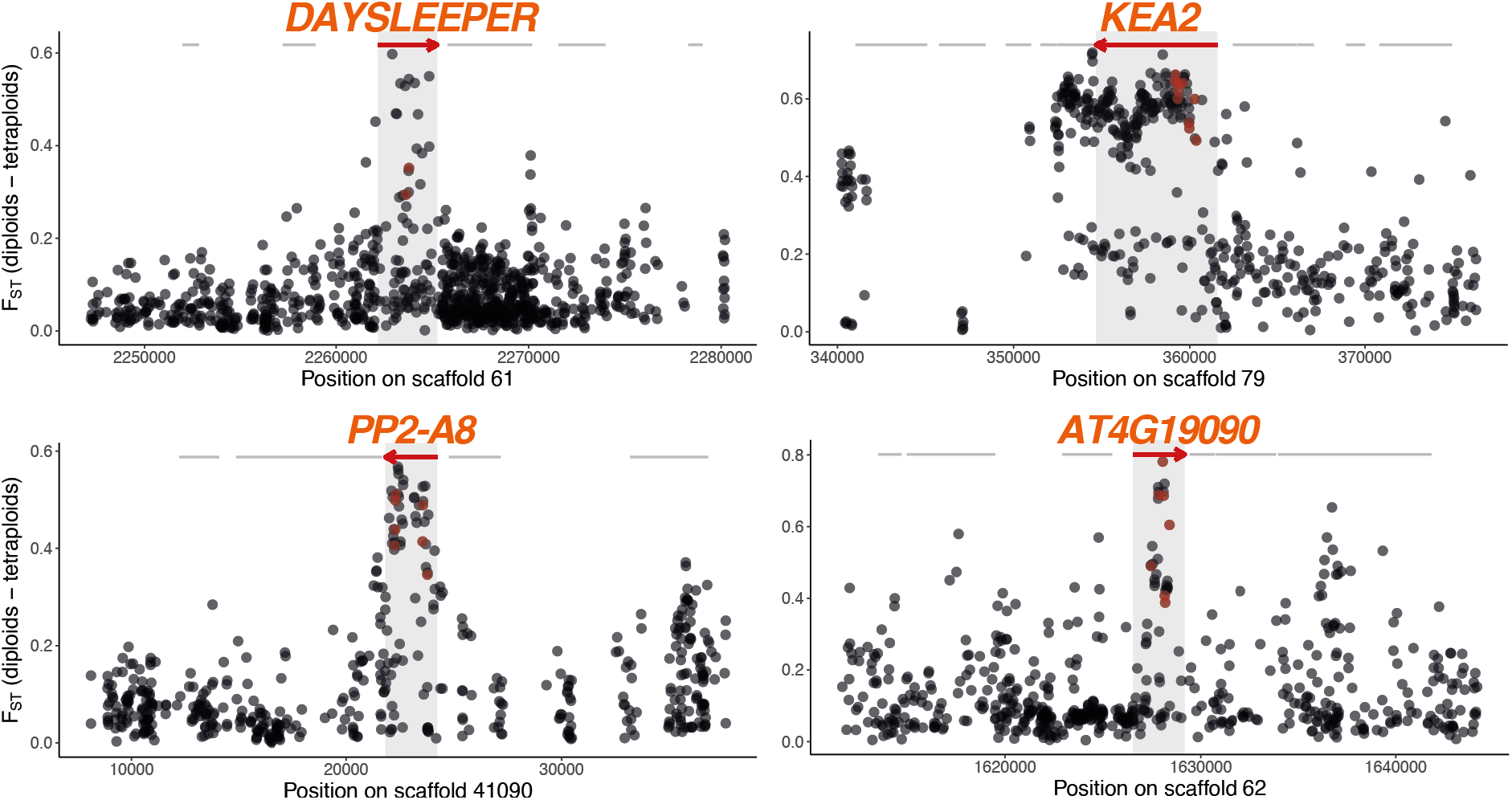
Selective sweep signatures at DNA management and ion homeostasis loci. Examples of selective sweep signatures among four loci (red) among Fst candidate genes. X-axis gives scaffold position in base pairs. Y-axis gives Fst values at single-nucleotide polymorphisms (dots) between diploid and autotetraploid *C. amara*. Red dots indicate fineMAV outlier SNPs. Red arrows indicate gene models overlapping top 1% Fst windows and grey lines indicate neighbouring gene coding loci.

### Evolution of stress, signalling, and ion homeostasis genes

The remainder of the enriched GO categories in *C. amara* revolved around a diversity of intracellular processes, including abiotic and biotic stress response, protein phosphorylation, root development, ABA signalling, and ion homeostasis. The intersection of these processes was often represented by several genes. For example, two of the top 20 highest-scoring SNPs in the genome-wide fineMAV analysis reside in SNF1-related protein kinase SnRK2.9 (*SI Appendix*, Table S5). SnRKs have been implicated in osmotic stress and root development (40, 41), and their activity also mediates the prominent roles of Clade A protein phosphatase 2C proteins in ABA and stress signalling (42). Interesting in this respect is a strong signature of selection in *HIGHLY ABA-INDUCED PP2C GENE 1*, a clade A *PP2C* protein (*SI Appendix*, Table S3). Stress-related phosphoinositide phosphatases are represented by *SAC9*, mutants of which exhibit constitutive stress responses (43). Diverse other genes related to these categories exhibit the strongest signatures of selection, such as *PP2-A8* (44) and *AT4G19090*, a transmembrane protein strongly expressed in young buds (45) (Fig. 2).

Given the observed increase in potassium and dehydration stress tolerance in first generation autotetraploid *A. thaliana* (5), it is very interesting that our window-based outliers include an especially dramatic selective sweep at *K^+^ Efflux Antiporter 2* (*KEA2*, Fig. 2), a K^+^ antiporter that modulates osmoregulation, ion, and pH homeostasis (46). Recent evidence indicates that *KEA2* is important for eliciting a rapid hyperosmotic-induced Ca^2+^ response to water limitation imposed by osmotic stress (47). The *KEA2* locus in autotetraploid *C. amara* features an exceptional ten fineMAV-outlier SNPs (Fig. 2, *SI Appendix*, Table S3, S5), indicating that the sweep contains a run of radical amino acid changes at high allele frequency difference between the ploidies, strongly suggesting a ploidy-selected functional change. We also detect *cation-chloride co-transporter 1* (*HAP 5*) a Na+, K+, Cl− co-transporter, involved in diverse developmental processes and Cl− homeostasis (48).

### Limited gene-level convergence between *C. amara* and *A. arenosa*

We hypothesized that WGD imposed strong, specific selection pressures leading to convergent directional selection on the same genes or at least on different genes playing a role in the same process (gene- or function-level convergence, respectively) between *C. amara* and *A. arenosa*. To test for this, we complemented our *C. amara* genome scan with an analysis of *A. arenosa* divergence outliers based on an expanded sampling relative to the original *A. arenosa* genome scan studies (13, 14). We selected the 80 diploid and 40 autotetraploid individuals sequenced most deeply in a recent range-wide survey (8) of genomic variation in *A. arenosa* (mean coverage depth per individual = 18; 160 haploid genomes sampled of each ploidy), and scanned for Fst outliers in 1 kb windows, as we did for *C. amara*. We identified 696 windows among 1% Fst outliers, overlapping 452 gene-coding loci (*SI Appendix*, Table S6), recovering results similar to (14), including the interacting set of 8 loci that govern meiotic chromosome crossovers. However, from this entire list of 452 *A. arenosa* WGD adaptation candidates, only six orthologous loci were shared with our 229 *C. amara* WGD adaptation candidates (Table 2). This degree of overlap was not significant (p = 0.42, Fisher’s exact test), indicating no excess convergence at the level of orthologous genes beyond random overlap. Similarly, there was no excess overlap among genes which harbour at least one candidate fineMAV substitution (3 overlapping candidate genes out of 120 in *C. amara* and 303 in *A. arenosa*; p = 0.27, Fisher’s exact test). This lack of convergence at the ortholog level may come as a surprise given the expected shared physiological challenges attendant to WGD (3).

**Table 2.**
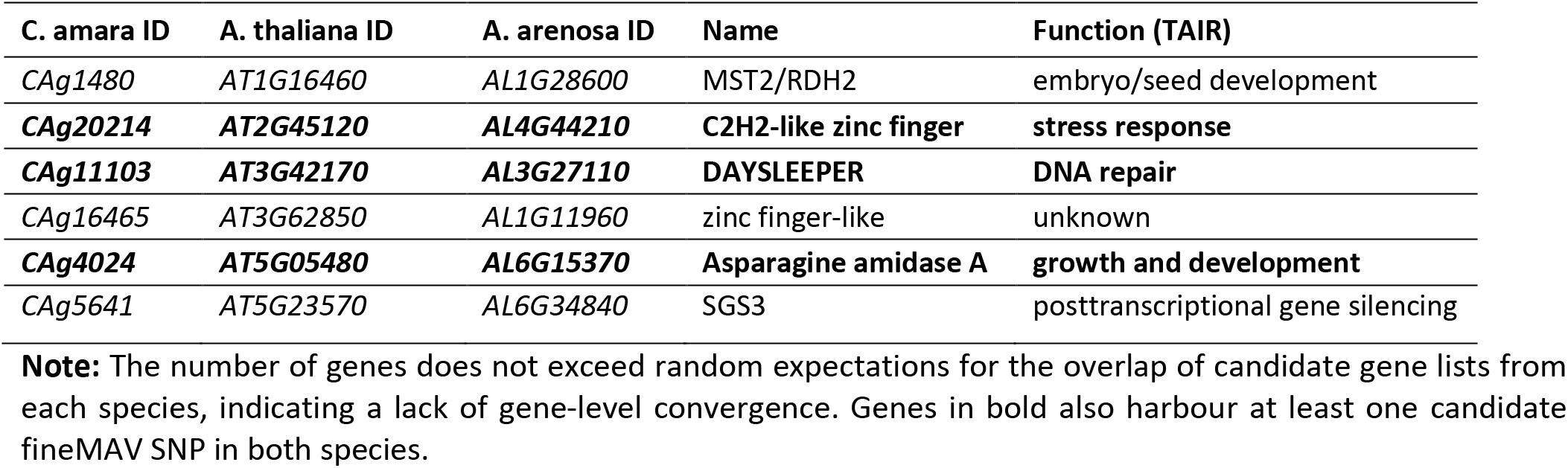
WGD adaptation candidates in both *A. arenosa* and *C. amara*

To determine whether we may have failed to detect convergent loci due to missing data or if genes that stand as top outliers in *A. arenosa* had few, but potentially functionally-implicated, differentiated SNPs in *C. amara*, we performed a targeted search in *C. amara* for the interacting set of meiosis proteins found to exhibit the most robust signatures of selection in *A. arenosa* (14). All meiosis-related orthologs in *C. amara* that also exhibit selection signatures in *A. arenosa* (13 in total) passed our data quality criteria and were included in our analyses. Three showed any signal at all by fineMAV analysis: *PDS5b* harbours an unusually high three fineMAV outlier SNPs, although it is not a window based Fst outlier. *ASY3*, which controls crossover distribution at meiosis, has only one fineMAV 1% outlier polymorphism.

### Error! Bookmark not defined

Finally, a regulator of endoreduplication, *CYCA2;3*, also harbours a single fineMAV 1% outlier SNP in *C. amara*, although it was not included in the Fst window analysis (number of SNPs < 20). However, upon inspection of Fst values of the (unusually low) 7 SNPs in the window overlapping this gene, the selection signal in CYCA2;3 would be high (mean Fst = 0.55). Thus, while we detect varying signal in these three meiosis-related genes following WGD, we do not see signals of selection in the conspicuous set of interacting crossover-controlling genes that were obvious in *A. arenosa* (14).

### Meiotic stability in *C. amara*

Despite our broad overall analysis of selection in *C. amara*, as well as a targeted assessment of the particular meiosis genes, we did not detect a signal of selection in those genes in *C. amara* (*SI Appendix*, Table S8). The *C. amara* autotetraploid is well-established lineage that underwent significant niche expansion in nature (18), but we still wondered if a contrast in meiotic behaviour underlies this difference in specific loci under selection. Therefore, we cytologically assessed the degree of male meiotic stability in *C. amara* (Fig. 3A). This revealed a low degree of stability in both *C. amara* cytotypes (mean proportion stable metaphase I cells in diploid maternal seed lines = 0.38 – 0.69, n = 133 scored cells; in tetraploids = 0.03 – 0.38; n = 348 scored cells; *SI Appendix*, Table S9). However, the degree of meiotic stability was lower in autotetraploids compared to diploids (differing proportion of stable to unstable meiotic cells for each ploidy; D = 62.7, df = 1, p < 0.0001, GLM with binomial errors; Fig. 3B, *SI Appendix*, Table S9), which corresponds with the lack of selection signal in crossover-controlling meiosis genes. Interestingly, we did find that the degree of stability was variable within each cytotype, suggesting the existence of standing genetic variation that controls stability. In contrast, higher frequencies of stable metaphase I cells (>80%) have been observed for diploid and autotetraploid *A. arenosa* (9). This, together with the observation of frequent clonal spreading of *C. amara* (49), indicates that the species has an ability to maintain stable populations, even under varying efficiencies of euploid gamete production, thus perhaps decreasing the urgency to fully stabilise meiosis in either cytotype. This, in turn, may have facilitated the establishment of the autotetraploid cytotype.

**Figure 3.**
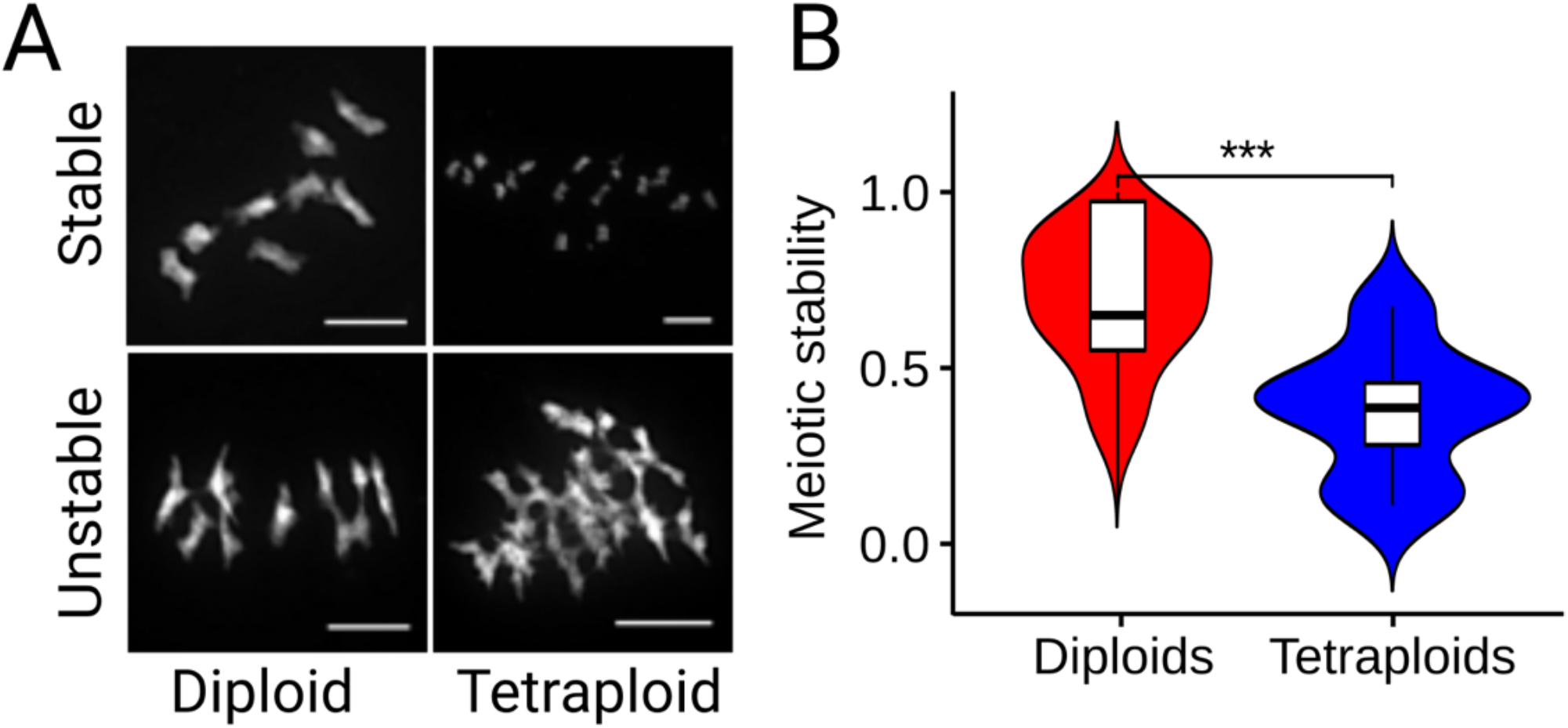
Variable meiotic stability in *C. amara*. **A,** An example of stable and unstable diploid (8 bivalents) and autotetraploid (16 bivalents) DAPI-stained meiotic chromosomes (diakinesis and metaphase I). Unstable meiosis is characterised by multivalent formation and interchromosomal connections. Scale bar corresponds to 10 μm. For a complete overview of all scored chromosome spreads see Figure S4. **B,** Distribution of meiotic stability (calculated as proportion of stable and partly stable to all scored meiotic spreads) in diploid and autotetraploid individuals of *C. amara.* *** - p < 0.001, GLM with binomial errors.

### Evidence for process-level convergence

While we found no excess convergence at the level of orthologous genes under selection, we speculated that convergence may occur nevertheless at the level of functional processes. To test this, we used two complementary approaches: overlap of GO term enrichment and evidence of shared protein function from interaction networks. First, of the 73 significantly (p < 0.05) enriched GO terms in *A. arenosa* (*SI Appendix*, Table S7), we found that five were identical to those significantly enriched in *C. amara*, which is more than expected by chance (p < 0.001, Fisher’s exact test; Table 3). In addition, some processes were found in both species, but were represented by slightly different terms, especially in the case of meiosis (“meiotic cell cycle” in *C. amara*, “meiotic cell cycle process” in *A. arenosa*: Tables S4 and S7). Remarkably, the relative ranking of enrichments of all five convergent terms was identical in both *C. amara* and *A. arenosa* (Table 3). This stands in strong contrast to the fact that *A. arenosa* presented an obvious set of physically and functionally interacting genes in the top two categories (‘DNA metabolic process’ and ‘chromosome organisation’), while the genes in these categories in *C. amara* are implicated in more diverse DNA management roles.

**Table 3.** Convergent processes under selection in both *C. amara* and *A. arenosa* following WGD.

| GO ID | Term | p-value<br>(*C. amara*) | p-value<br>(*A. arenosa*) | Enrichment<br>(*C. amara*) | Enrichment<br>(*A. arenosa*) |
| --- | --- | --- | --- | --- | --- |
| GO:0006259 | DNA metabolic process | 6.50E-08 | 8.20E-04 | 3.72 | 2.46 |
| GO:0051276 | chromosome organization | 0.019 | 2.10E-04 | 1.98 | 2.01 |
| GO:0009738 | abscisic acid-activated signalling pathway | 0.032 | 0.022 | 2.54 | 2.10 |
| GO:0071215 | cellular response to abscisic acid stimulation | 0.048 | 0.04 | 2.30 | 1.90 |
| GO:0097306 | cellular response to alcohol | 0.048 | 0.04 | 2.30 | 1.90 |

**Note:** p-values given are Fisher's exact test using the conservative 'elim' method, which tests for enrichment of terms from the bottom of the GO hierarchy to the top and discards any genes that are significantly enriched in a descendant GO terms (Methods). 'Enrichment' refers to fold enrichment.

Second, we sought for evidence that genes under selection in *C. amara* might interact with those found under selection in *A. arenosa*, which would further support process-level convergence between the species. Thus, we took advantage of protein interaction information from the STRING database, which provides an estimate of proteins’ joint contributions to a shared function (50). For each *C. amara* WGD adaptation candidate we searched for the presence of STRING interactors among the *A. arenosa* WGD adaptation candidates, reasoning that finding such an association between candidates in two species may suggest that directional selection has targeted the same processes in both species through different genes. Following this approach, we found that out of the 229 *C. amara* WGD adaptation candidates, 90 were predicted to interact with at least one of the 452 WGD adaptation candidates in *A. arenosa*. In fact, 57 likely interacted with more than one *A. arenosa* candidate protein (Fig. 4 and *SI Appendix*, Table S10). This level of overlap was greater than expected by chance (p = 0.001 for both “any interaction” and “more-than-one interaction”, as determined by permutation tests with the same database and 1000 randomly generated candidate lists).

**Figure 4.**
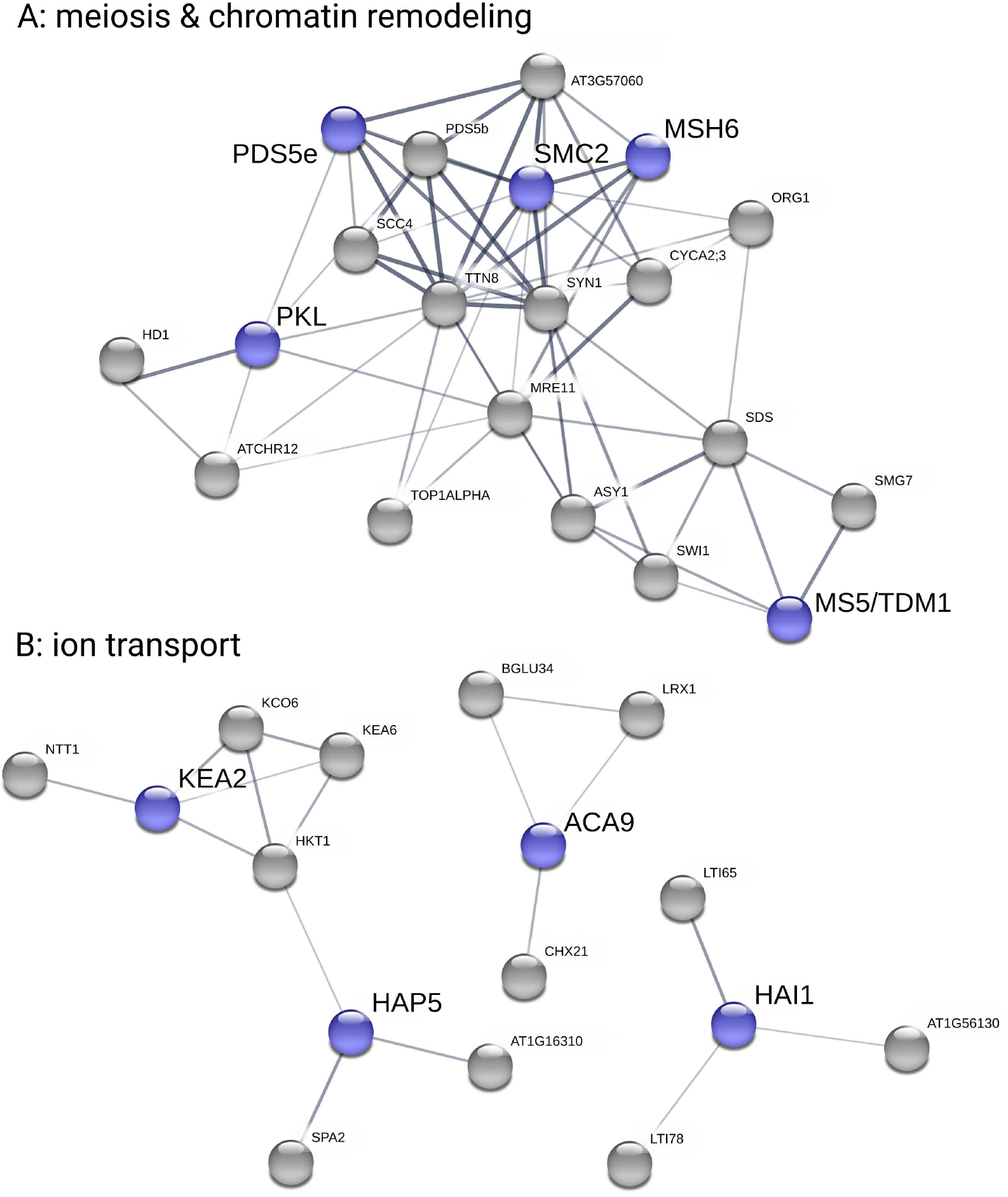
Evidence for functional parallelism between *C. amara* and *A. arenosa* following independent WGDs. Plots show *C. amara* candidate genes in blue and STRING-associated *A. arenosa* candidate genes in grey. We used only medium confidence associations and higher (increasing thickness of lines connecting genes indicates greater confidence). **A**, meiosis- and chromatin remodelling-related genes. **B**, ion transport-related genes.

Several large STRING clusters were evident among WGD adaptation candidates in *C. amara* and *A. arenosa* (Fig. 4). The largest of these clusters centre on genome maintenance, specifically meiosis and chromatin remodelling (Fig 4A), and ion homeostasis (especially K+ and Ca2+), along with stress (ABA) signalling (Fig 4B), consistent with the results of GO analysis. Taken together, both STRING and GO analyses support our hypothesis of functional convergence of these processes following WGD in *C. amara* and *A. arenosa.*

## Conclusion

Given the expected shared challenges attendant to WGD in *C. amara* and *A. arenosa*, we hypothesised at least partially convergent evolutionary responses to WGD. While we found obvious convergent recruitment at the level of functional processes, we did not detect excess convergence at the genic level. This was consistent with the probable absence of shared standing variation between these species (51), which are 17 million years diverged. Nevertheless, we note that if any shared variation has persisted, it was not selected upon convergently in both young autotetraploids, thus strengthening the conclusion that the genes selected in response to WGD are not highly constrained between these two species. The most prominent difference we observed here is the lack of an obvious coordinated evolutionary response in genes stabilizing early meiotic chromosome segregation in *C. amara*, relative to the striking coevolution of physically and functionally interacting proteins governing crossover formation in *A. arenosa*. This could be explained to some extent by the observation that in *C. amara* both diploids and autotetraploids are not uniformly meiotically stable, and autotetraploids may therefore enjoy a less strict reliance on the generation of a high percentage of euploid gametes. This may allow the decoupling of crossover reduction from broader changes across meiosis and other processes we observe. This is not to say that we see no signal of WGD adaptation in *C. amara*: factors governing timing during later meiosis, especially the exit from meiotic divisions as evidenced by the interacting trio of *SMG7, SDS* and *MS5*, along with other chromatin remodelling factors and DNA repair-related proteins, such as *MSH6* and *DAYSLEEPER* give very strong signals. The convergent functions we did detect (other meiosis processes, chromosome organisation/chromatin remodelling, ABA signalling and ion transport) provide first insights into the salient challenges associated with WGD.

We conclude that evolutionary solutions to WGD-associated challenges vary from case to case, suggesting minimal constraint. This may explain how many species manage to thrive following WGD and, once established as polyploids, experience evolutionary success. In fact, we envision that the meiotic instability experienced by some WGD lineages, such as *C. amara*, could serve as a diversity-generating engine promoting large effect genomic structural variation, as has been observed in aggressive polyploid gliomas (3).

## Materials and Methods

### Reference Genome Assembly and Alignment

We generated a *de novo* assembly using the 10x Genomics Chromium approach. In brief, a single diploid individual from pop LUZ (*SI Appendix*, Table S11) was used to generate a single Chromium library, sequenced using 250PE mode on an Illumina sequencer, and assembled with Supernova version 2.0.0. This assembly had an overall scaffold N50 of 1.82mb. An assessment of genome completeness using BUSCO (version 3.0.2) (52) for the 2,251 contigs >=10kb was estimated at 94.8% (1365/1440 BUSCO groups; *SI Appendix*, Table S12).

#### BioNano Plant Extraction protocol

High molecular weight DNA extraction and 10x synthetic long read library construction detailed methods are provided in *SI Appendix*, Methods. The libraries were run at 250 paired end format on an Illumina HiSeq.

#### Sequencing and assembly and assembly QC

Raw reads were subsampled to 90 M reads and assembled with Supernova 2.0.0 (10x Genomics), giving a raw coverage of 60.30x and an effective coverage of 47.43x. The estimated molecule length was 44.15 kb. The assembly size, counting only scaffolds longer than 10kb was 159.53 Mb and the Scaffold N50 was 1.82MB. The k-mer estimate for the genome size was 225.39 MB, hence we are missing 16.61% from the assembly by retaining only contigs longer than 10Kb. We further scaffolded the assembly using the published *Cardamine hirsuta* genome using *graphAlign* (53) and *Nucmer* (54).

#### Gene Calling and Annotation

The ‘plants set’ database ‘embryophyta_odb9.tar.gz’ was downloaded from http://busco.ezlab.org/ and used to assess orthologue presence/absence in our *C. amara* genome annotation. Running BUSCO gave Augustus (55) results via BUSCO HMMs to infer where genes lie in the assembly and returning their protein sequences. A blast (v. 2.2.4) database was built for Brassicales (taxid: 3699) by downloading ~ 1.26M protein sequences from https://www.ncbi.nlm.nih.gov/taxonomy/ and the Augustus-predicted proteins were annotated via Interproscan (56) and blast2go (57).

#### Orthogrouping and Reciprocal Best Blast Hits

We performed an orthogroup analysis using Orthofinder version 2.3.3 (58). to infer orthologous groups (OGs) from four species (*C. amara, A. lyrata, A. thaliana, C. pyrenaica*). A total of 21,618 OGs were found. Best reciprocal blast hits (RBHs) for *C. amara* and *A. thaliana* genes were found using BLAST version 2.9.0. *C. amara* genes were then assigned an *A. thaliana* gene ID for GO enrichment analysis via the following protocol: First, if the genes’ OG contained only one *A. thaliana* gene ID, that gene ID was used. If the OG contained more than one *A. thaliana* gene, then the RBH was taken. If there was no RBH the *A. thaliana* gene ID, then the OG gene with the lowest E-value in a BLAST versus the TAIR10 database was taken. If no OG contained the *C. amara* gene, then the RBH was taken. Finally, if there was no OG or RBH then the gene with the lowest E-value in a BLAST versus the TAIR10 database was taken. BLASTs versus the TAIR10 database were performed during December 2019.

### Population resequencing and genome scans for selection

#### Sampling design

To isolate genomic regions subjected to directional selection acting specifically between diploids and autotetraploids, we sampled a set of two diploid and two tetraploid populations (Fig. 1C). We used comparisons between populations of the same ploidy to constitute a null model for shared heterogeneity in genetic differentiation arising through processes unrelated to WGD (following an approach successfully applied in (21)).

#### Library preparation and sequencing

We extracted DNA in triplicate from 25 individuals for each of the following populations: CEZ (4x), PIC (4x), VKR (2x), and LUZ (2x). All plants used for DNA extraction were verified for expected ploidy by flow cytometry. We then pooled samples of each population, constructed Illumina Truseq libraries (Illumina), and sequenced them on an Illumina NextSeq at a 150 base pair, paired-end specification.

#### Data preparation, alignment, and genotyping

Fastq files from the two runs were combined and concatenated to give an average of 30.5 million reads per sample. Data processing steps are given in *SI Appendix*, Methods, along with variant calling steps.

#### Population genetic structure

We first calculated genome-wide between-population metrics (Nei’s Fst (59) and allele frequency difference). We calculated allele frequencies (AF) as the average AF of all the pools. The AF in individual pools has been calculated as the fraction of the total number of reads supporting the alternative allele (60). We used the python3 PoolSeqBPM pipeline, designed to input pooled data (https://github.com/mbohutinska/PoolSeqBPM). Then we inferred relationships between populations as genetic distances calculated over putatively neutral four-fold degenerate sites using principal component analysis (PCA) implemented in *adegenet* (61).

#### Window-based selection scan using a quartet design

We performed a window-based Fst (59) scan for directional selection in *C. amara*, taking advantage of quartet of two diploid and two autotetraploid populations (Fig. 1C). Using such quartet design, we identified top candidate windows for selective sweeps associated with ploidy differentiation, while excluding differentiation patterns private to a single population or ploidy-uninformative selective sweeps. To do so, we calculated Fst for 1 kb windows with minimum 20 SNPs for all six population pairs in the quartet (Fig. 1C) and ranked windows based on their Fst value. We excluded windows which were top 1% outliers in diploid-diploid (dd in Fig. 1C) or autotetraploid-autotetraploid (tt) populations contrasts, as they represent variation inconsistent with diploid-autotetraploid divergence but rather signal local differentiation within a cytotype. Next, we assigned ranks to each window based on the Fst values in four diploid-autotetraploid contrasts and identified windows being top 1% outliers of minimum rank sum.

To account for possible confounding effect of comparing windows from genic and non-genic regions, we calculated the number of base pairs overlapping with any gene within each window. There was not any relationship between the proportion of genic space within a window and Fst (Pearson’s R^2^ = −0.057, Figure S3), indicating that our analyses were unaffected by unequal proportion of genic space in a window.

In *A. arenosa*, we performed window-based Fst scan for directional selection using the same criteria as for *C. amara* (1kb windows, min 20 SNPs per window). We did not use the quartet design as the range-wide dataset of 80 diploid and 40 autotetraploid individuals drawn from many populations assured power to detect genomic regions with WGD-associated differentiation.

#### FineMAV

We adopted the Fine-Mapping of Adaptive Variation (fineMAV(22)), and modified it to fit the resources available for *r*eference genome of *C. hirsuta*. Specifically, we replaced CADD, the functional score available for amino acids in human reference (22, 62), by the Grantham score (63), which is a purely theoretical amino acid substitution value, encoded in the Grantham matrix, where each element shows the differences of physicochemical properties between two amino acids. Details on FineMAV data processing are given in *SI Appendix*, Methods.

#### Arabidopsis arenosa population genomic dataset

We complemented our analysis of adaptation to WGD in *C. amara* with analysis of *A. arenosa*, based on an expanded sampling (8) relative to the original *A. arenosa* WGD adaptation studies (13, 14). We first aligned the short read sequences to the *A. lyrata* reference genome, called variants and filtered as previously using the Genome Analysis Toolkit (GATK 3.5 and 3.6 (64)). We used a subset of the dataset consisting of 80 diploid individuals (samples selected based on the highest mean depth of coverage) and 40 tetraploid individuals from populations unaffected by secondary introgression from diploid lineages (8). Such sub-sampling gave us a balanced number of 160 high-quality haploid genomes of each ploidy suitable for selection scans. Finally, we filtered each subsampled dataset for genotype read depth > 8 and maximum fraction of missing genotypes < 0.5 in each lineage. We calculated Fst using python3 ScanTools pipeline (github.com/mbohutinska/ScanTools_ProtEvol). All subsequent analyses were performed following the same procedure as with *C. amara* data.

#### GO enrichment analysis

To infer functions significantly associated with directional selection following WGD, we performed gene ontology enrichment of gene list using the R package topGO (65), using *A. thaliana* orthologs of *C. amara/A. lyrata* genes, obtained using biomaRt (66). We used Fisher’s exact test with conservative ‘elim’ method, which tests for enrichment of terms from the bottom of the GO hierarchy to the top and discards any genes that are significantly enriched in a descendant GO terms (67). We used ‘biological process’ ontology with minimum node size 150 genes.

#### Protein associations from STRING database

We searched for association among *C. amara* and *A. arenosa* candidate genes using STRING (50) database. We used multiple proteins search in *A. thaliana*, with text mining, experiments, databases, co-expression, neighbourhood, gene fusion and co-occurrence as information sources. We used minimum confidence 0.4 and retained only 1st shell associations (proteins that are directly associated with the candidate protein: i.e., neighbouring circles in the network).

#### Quantifying convergence

We considered convergent candidates all candidate genes or significantly enriched GO categories that overlapped across both species. Convergent candidate genes had to be members of the same orthogroups (58). To test for higher than random number of overlapping items we used Fisher’s Exact Test for Count Data in R (68).

### Cytological assessment of meiotic stability

#### Chromosome preparation

Whole young inflorescences were fixed in freshly prepared ethanol:acetic acid (3:1) overnight, transferred into 70 % ethanol and later stored at −20 °C until use. Meiotic chromosome spreads were prepared from anthers according to (69). Briefly, after washing in citrate buffer (10 mM sodium citrate, pH 4.8), selected flower buds were digested using a 0.3 % mix of pectolytic enzymes (cellulase, cytohelisase, pectolyase; Sigma-Aldrich Corp., St. Louis, MO) in citrate buffer for c. 3 h. Individual anthers were dissected and spread in 20 μl of 60 % acetic acid on a microscope slide placed on a metal hot plate (50 °C), fixed by ethanol:acetic acid (3:1) and the preparation was dried using a hair dryer. SuiSI Table Slides were postfixed in freshly prepared 4 % formaldehyde in distilled water for 10 min and air-dried. The preparations were stained with 4ʹ,6-diamidino-2-phenylindole (DAPI; 2 μg/ml) in Vectashield (Vector Laboratories, Peterborough, UK). Fluorescence signals were analysed using an Axioimager Z2 epifluorescence microscope (Zeiss, Oberkochen, Germany) and CoolCube CCD camera (MetaSystems, Newton, MA).

#### Meiotic stability scoring

In diploids, chromosome spreads with 8 bivalents were scored as “stable meiosis”, 7-6 as “partly stable”, 5-4 as “partly unstable”, and <4 as “unstable”. In autotetraploids, chromosome spreads with 16 bivalents were scored as “stable meiosis”, 14-12 as “partly stable”, 10-8 as “partly unstable”, and <8 as “unstable”. We report a mean value of meiotic stability for each ploidy calculated over “stable meiosis” and over sum of “stable meiosis” and “partly stable” categories. Difference in meiotic stability between diploids and autotetraploids (Fig. 3B) is reported for the sum of “stable” and “partly stable” categories. However, considering only “stable meiosis” category does not qualitatively affect the results (i.e. the degree of meiotic stability is significantly lower in tetraploids, D = 125.7, df = 1, p < 0.0001, GLM with binomial errors). Photos of all spreads scored are shown in Figure S4.

## Acknowledgements

The authors thank Veronika Konečná for assistance with the map figure and Doubravka Požárová and Paolo Bartolić for help with field collections. This work was supported by the European Research Council (ERC) under the European Union’s Horizon 2020 research and innovation programme [grant number ERC-StG 679056 HOTSPOT], via a grant to LY. Additional support was provided by Czech Science Foundation (project 20-22783S to FK, 19-03442S to TM and 19-06632S), by Charles University (project Primus/SCI/35 to FK), by the long-term research development project No. RVO 67985939 of the Czech Academy of Sciences and by the CEITEC 2020 project (grant LQ1601).

## Data Availability

Sequence data that support the findings of this study have been deposited in the Sequence Read Archive (SRA; https://www.ncbi.nlm.nih.gov/sra) with the study codes SRP156117 702 (*A. arenosa* data, released) and SRPXXXXXX (*C. amara* data, released upon publication). All script are available at github.com/mbohutinska/PoolSeqBPM (Fst-based selection scans and all following analyses) and github.com/paajanen/meiosis_protein_evolution (fineMAV scan).

## Author Contributions

LY conceived the study. MB, MA, PP, SB, TM and PM performed analyses. PM and TM performed laboratory experiments. PM, FK, SB, and MB performed field collections. LY and MB wrote the manuscript with input from all authors. All authors approved of the final manuscript.

## Materials & Correspondence

All requests should be addressed to Levi Yant at

## Supplemental Materials and Methods

### BioNano Plant Extraction protocol

Fresh young leaves of the *C. amara* accession LUZ were collected after 48-hour treatment in the dark. DNA was extracted by the Earlham Institute’s Platforms and Pipelines group following an IrysPrep “Fix‘n’Blend” Plant DNA extraction protocol supplied by BioNano Genomics. 2.5 g of fresh young leaves were fixed with 2% formaldehyde. After washing, leaves were disrupted and homogenized in the presence of an isolation buffer containing PVP10 and BME to prevent oxidation of polyphenols. Triton X-100 was added to facilitate the release of nuclei from the broken cells. The nuclei were then purified on a Percoll cushion. A nuclei phase was taken and washed several times in isolation buffer before embedding into low melting point agarose. Two plugs of 90 μl were cast using the CHEF Mammalian Genomic DNA Plug Kit (Bio-Rad 170-3591). Once set at 4°C the plugs were added to a lysis solution containing 200 μl proteinase K (QIAGEN 158920) and 2.5 ml of BioNano lysis buffer in a 50 ml conical tube. These were put at 50°C for 2 hours on a thermomixer, making a fresh proteinase K solution to incubate overnight. The 50 ml tubes were then removed from the thermomixer for 5 minutes before 50 μl RNAse A (Qiagen158924) was added and the tubes returned to the thermomixer for a further hour at 37°C. The plugs were then washed 7 times in Wash Buffer supplied in Chef kit and 7 times in 1xTE. One plug was removed and melted for 2 minutes at 70°C followed by 5 minutes at 43°C before adding 10 μl of 0.2 U /μl of GELase (Cambio Ltd G31200). After 45 minutes at 43°C the melted plug was dialysed on a 0.1 μM membrane (Millipore VCWP04700) sitting on 15 ml of 1xTE in a small petri dish. After 2 hours the sample was removed with a wide bore tip and mixed gently 5 times and left overnight at 4°C.

### 10X library construction

DNA material was diluted to 0.5ng/ul with EB (Qiagen) and checked with a QuBit Flourometer 2.0 (Invitrogen) using the QuBit dsDNA HS Assay kit (Table S11). The Chromium User Guide was followed as per the manufacturer’s instructions (10X Genomics, CG00043, Rev A). The final library was quantified using qPCR (KAPA Library Quant kit [Illumina] and ABI Prism qPCR Mix, Kapa Biosystems). Sizing of the library fragments was checked using a Bioanalyzer (High Sensitivity DNA Reagents, Agilent). Samples were pooled based on the molarities calculated using the two QC measurements. The library was clustered at 8 pM with a 1% spike in of PhiX library (Illumina). The pool was run on a HiSeq2500 150bp Rapid Run V2 mode (Illumina). The following run metrics were applied: Read 1: 250 cycles, Index 1: 8 cycles, Index 2: 0 cycles and Read 2: 250 cycles.

### Data preparation, alignment, and genotyping

Fastq files from the two runs were combined and concatenated to give an average of 30.5 million reads per sample. Adapter sequences were removed via the cutadapt software (version 1.9.1) (1) and quality trimmed via Sickle (version 33) (2) to generate only high-quality reads (Phred score >=30) of 30bp or more, resulting in an average of 27.9 million reads per sample. Using samtools (v. 1.7) (3) and bwa (v. 0.7.12) (4) software, the quality-filtered reads were aligned against two references: 89.3% of reads mapped to our *C. amara* assembly, while only 74.5% to *C. hirsuta*. We retained only the alignment to *C. amara* for all analysis. Using the picard software tool (v. 1.134) (5), first duplicate reads were removed via ‘MarkDuplicates’ followed by the addition of read group IDs to the bam files via ‘AddOrReplaceReadGroups’. Finally, to handle the presence of indels, GATK (v. 3.6.0) (6) was used to realign reads to the *C. amara* assembly via ‘RealignerTargetCreator’ and ‘IndelRealigner’.

### Variant Calling

Text files describing sample populations and ploidy were prepared, and variants called for the 12 bam files using Freebayes (v. 1.1.0.46)(7) to generate a single VCF output. Due to working with pooled (high ploidy) samples, Freebayes was run with ‘--pooled-discrete’ (assumes samples result from pooled sequencing). In addition, the software was restricted to biallelic sites (‘--use-best-n-alleles 2’) and indel sites were excluded (’--no-indels’). The VCF was filtered via bcftools (v 1.8) (8) to remove sites where the read depth was < 10 or greater than 1.6x the second mode (determined as 1.6 × 31 = 50, Figure S2).

### FineMAV

We downloaded coding sequences from the *C. hirsuta* genomic resources web site http://chi.mpipz.mpg.de/download/annotations/carhr38.cds.fa and mapped to *C. amara* using gmap. The resulting sam file was converted to bam-format, sorted and indexed via samtools (v. 1.7) (3), and then converted to GTF-format via the ‘convert’ script in Mikado (v1.2.3) (9) which was subsequently used to build a snpEFF (v. 4.3) (10) database. We estimated the population genetic component of fineMAV (see (11) for details on calculations) using allele frequency information at each site (considering minor frequency allele as derived) and DAP parameter of 3.5. Finally, for each amino acid substitution, we assigned Grantham scores, together with population genetic component of fineMAV, using a custom scripts in Python 2.7.10 and the Biopython 1.69 package. We identified the top 1% outliers and considered them the final candidates identified in fineMAV analysis. All the calculations were performed using code available at (github.com/paajanen/meiosis_protein_evolution).

## Supplemental Figures

**Figure S1:**
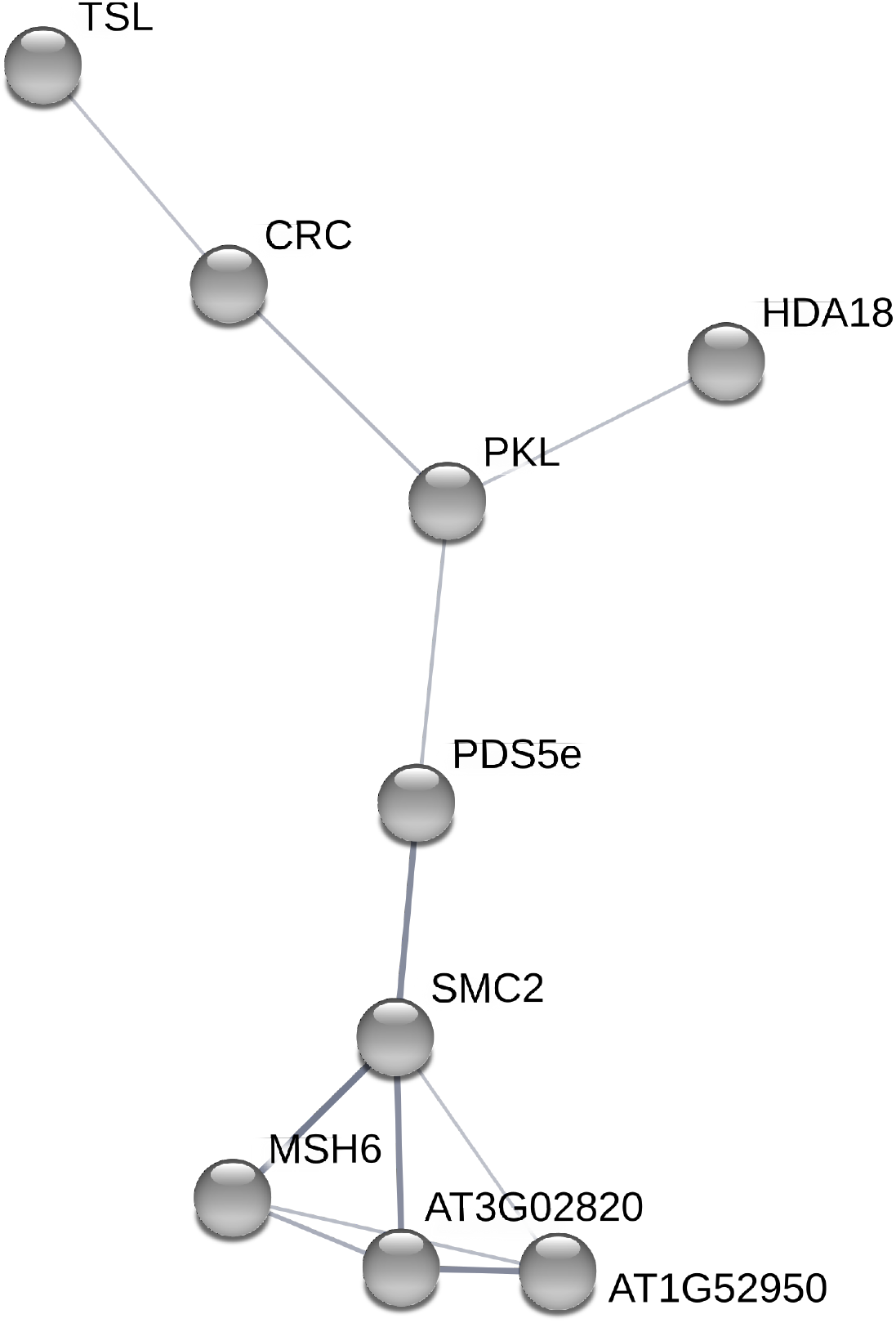
*C. amara* candidate meiosis gene associations as identified by STRING. We used only medium confidence associations and higher (shown as thickness of lines connecting genes).

**Figure S2:**
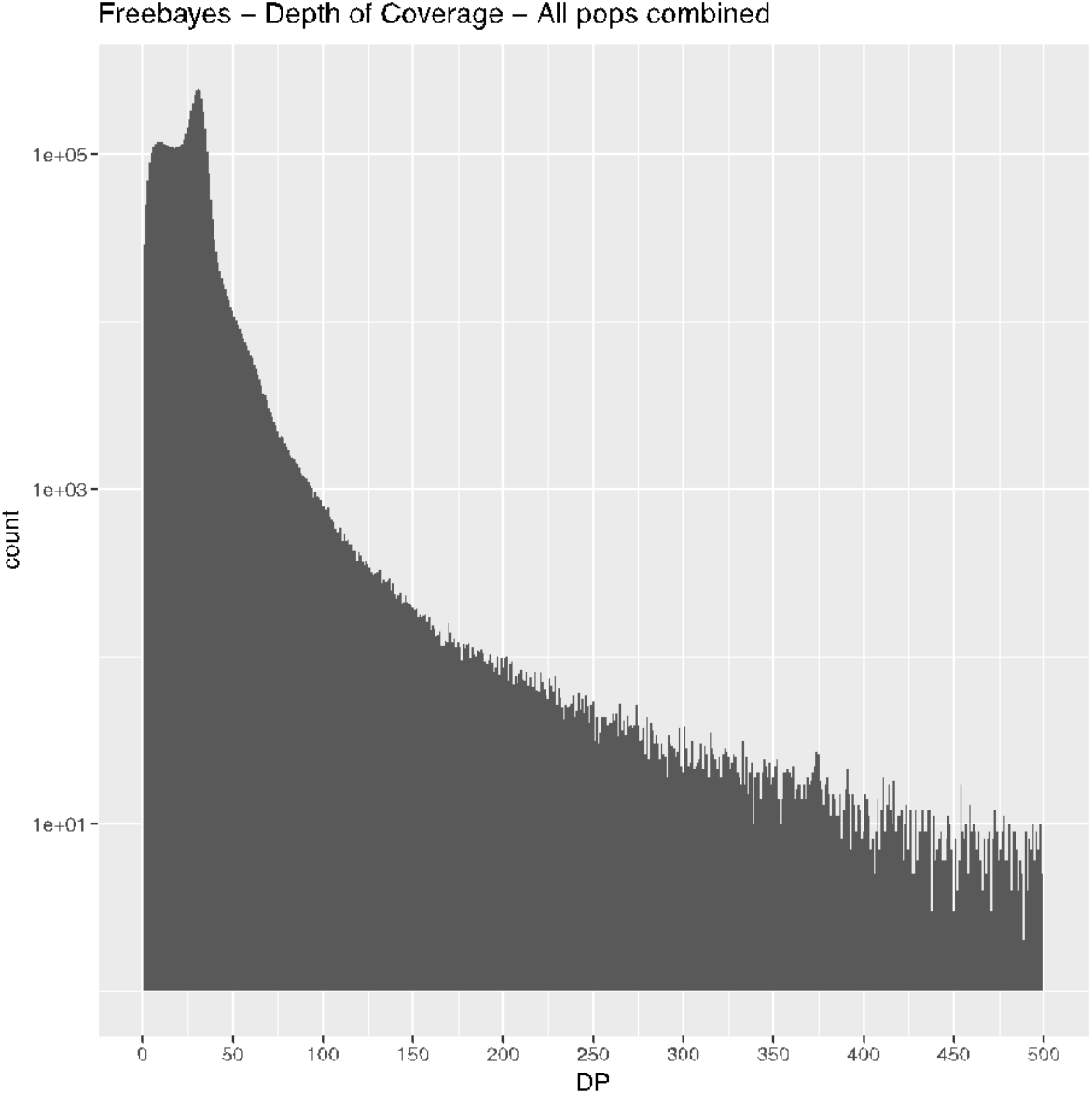
Distribution of read depth over all sequenced samples.

**Figure S3:**
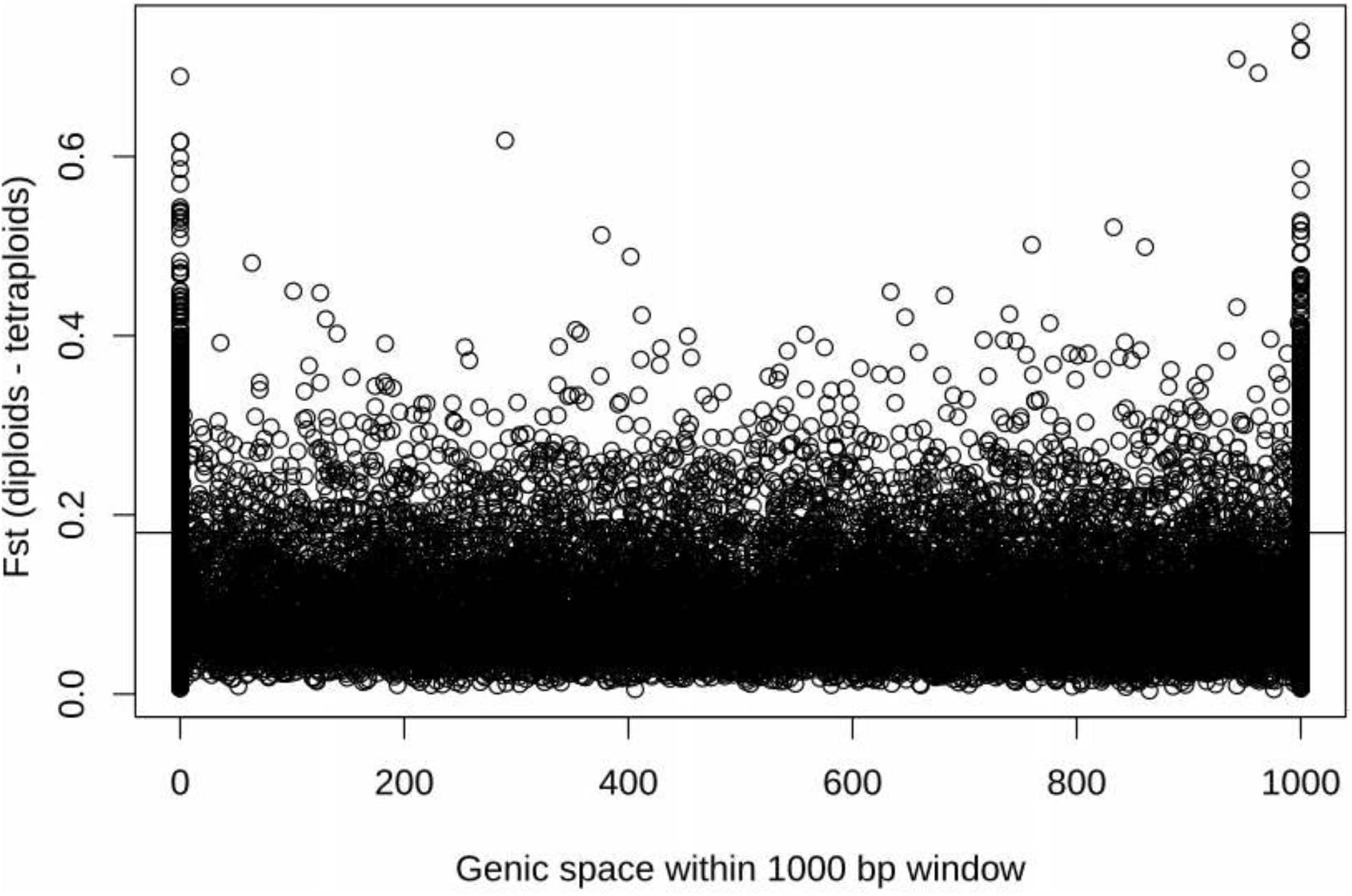
Relationship between the proportion of genic space within a window and Fst.

**Figure S4.**
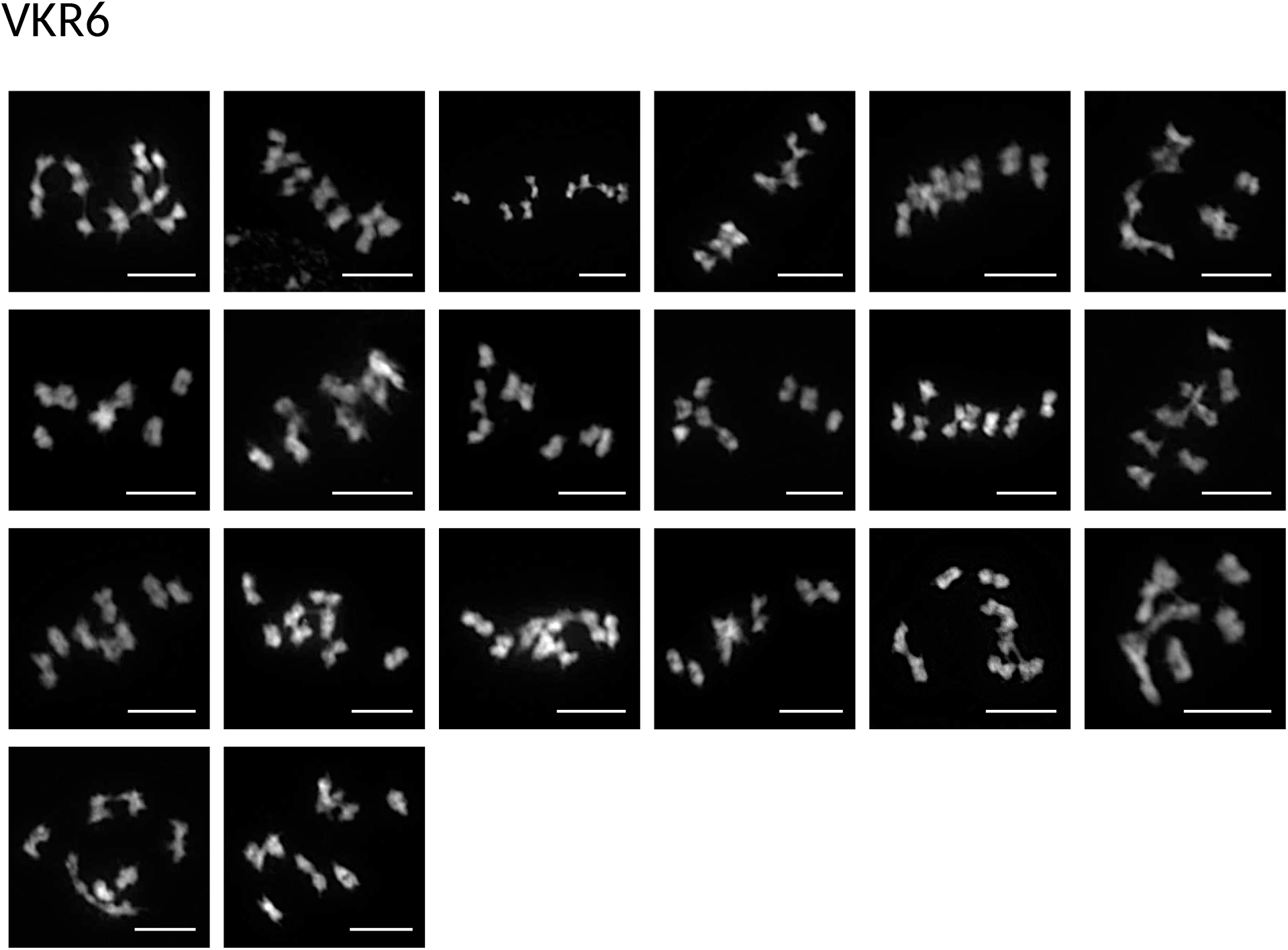
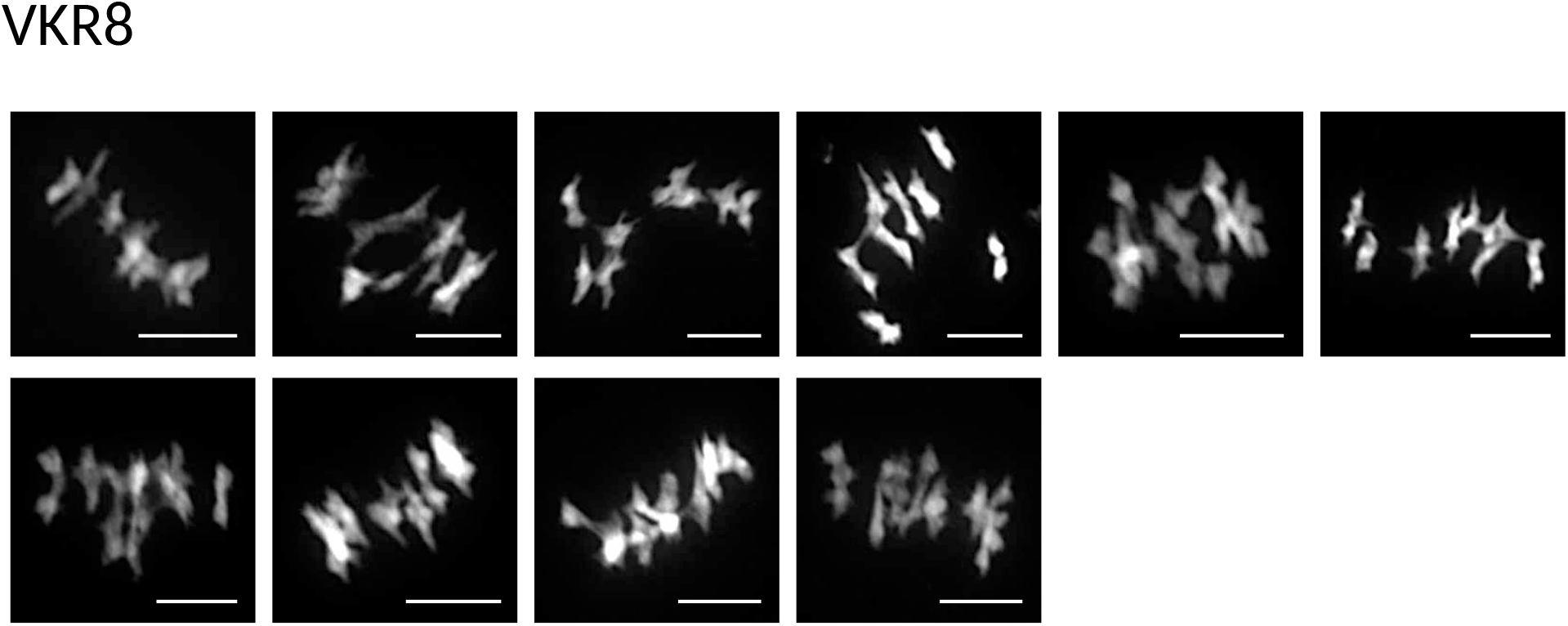
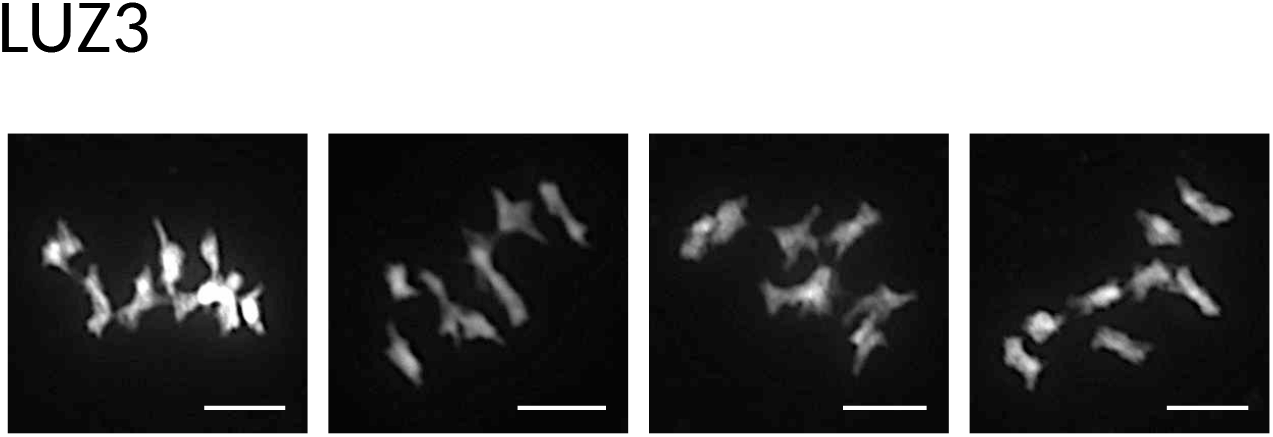
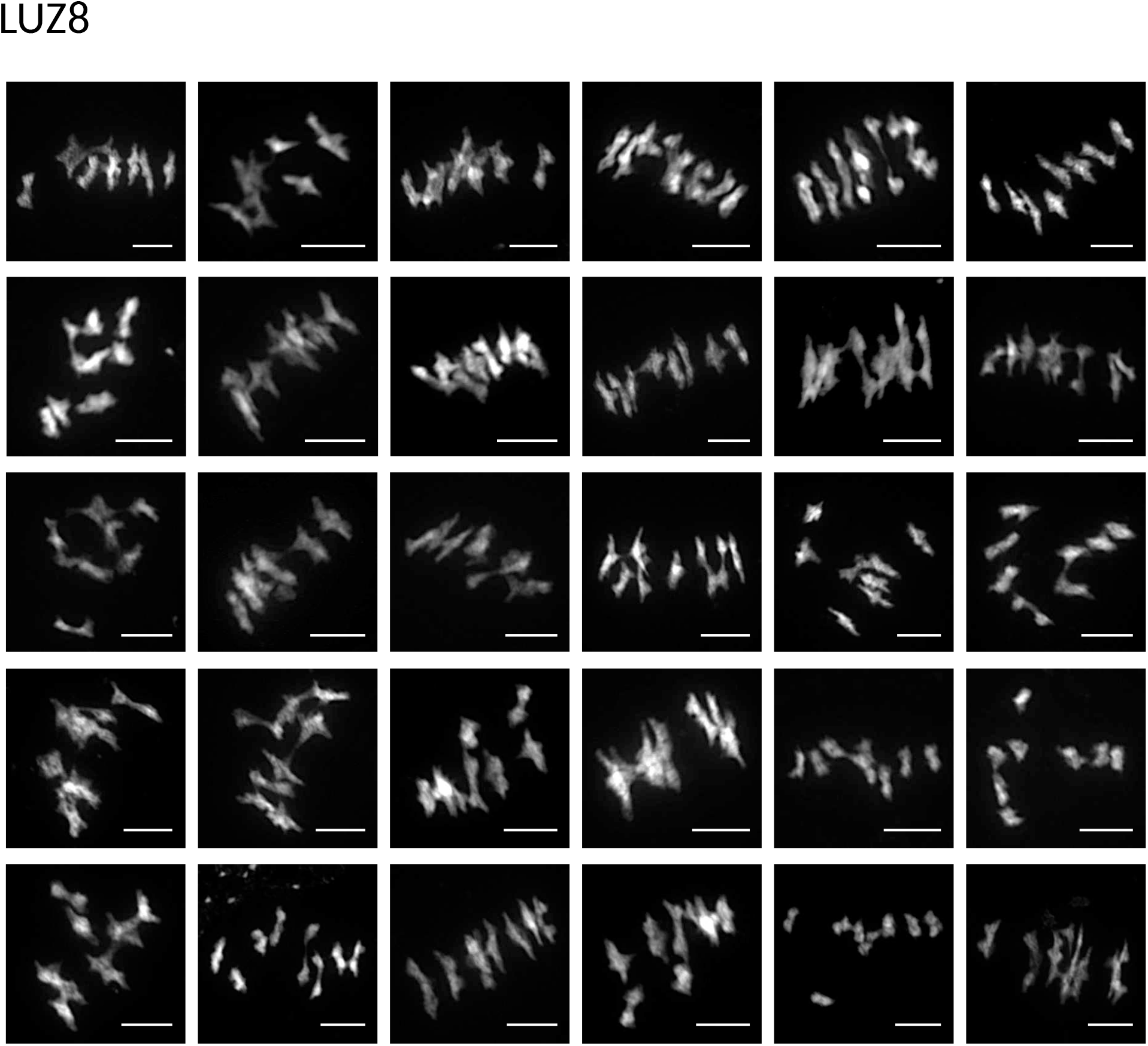
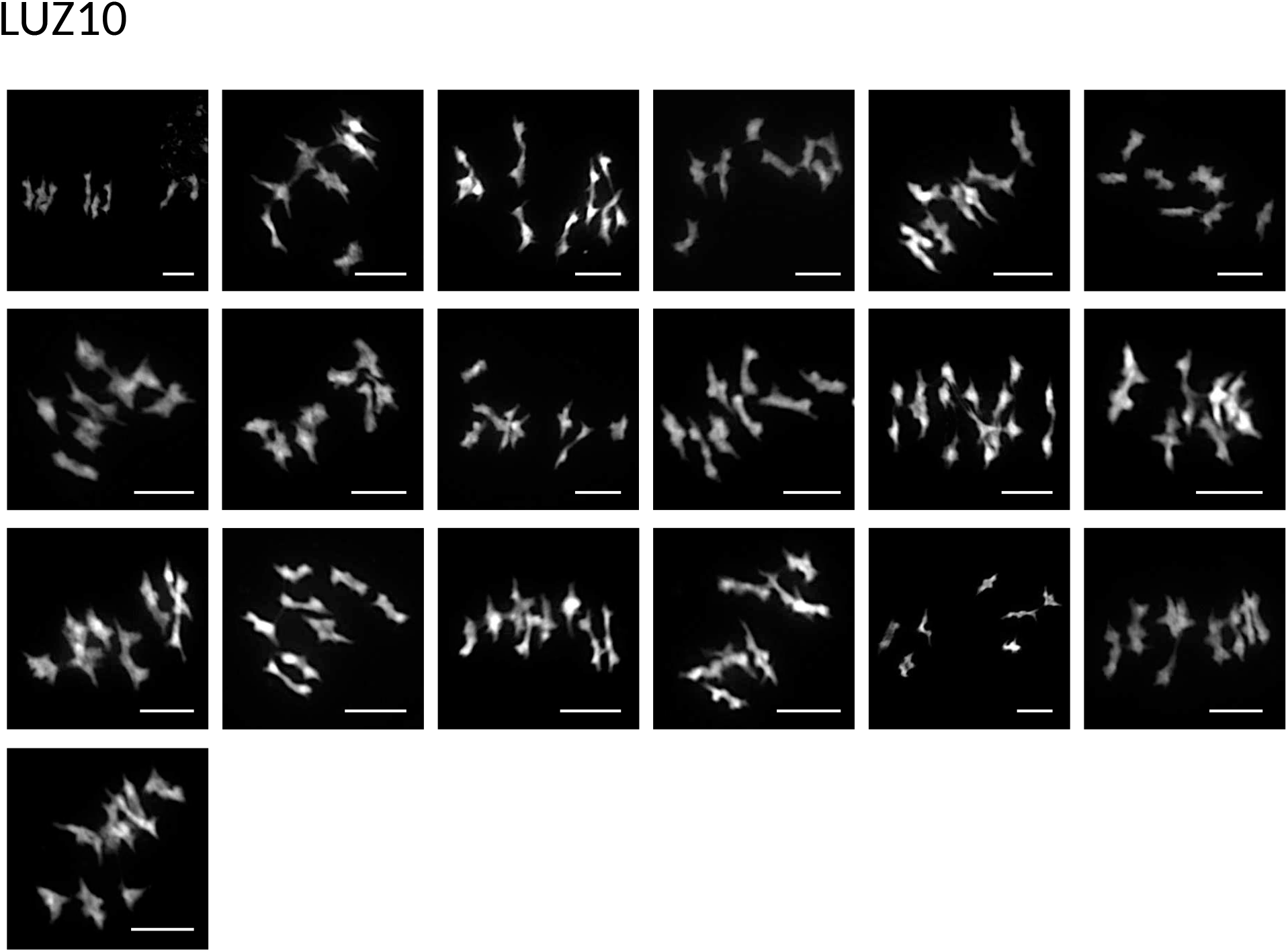
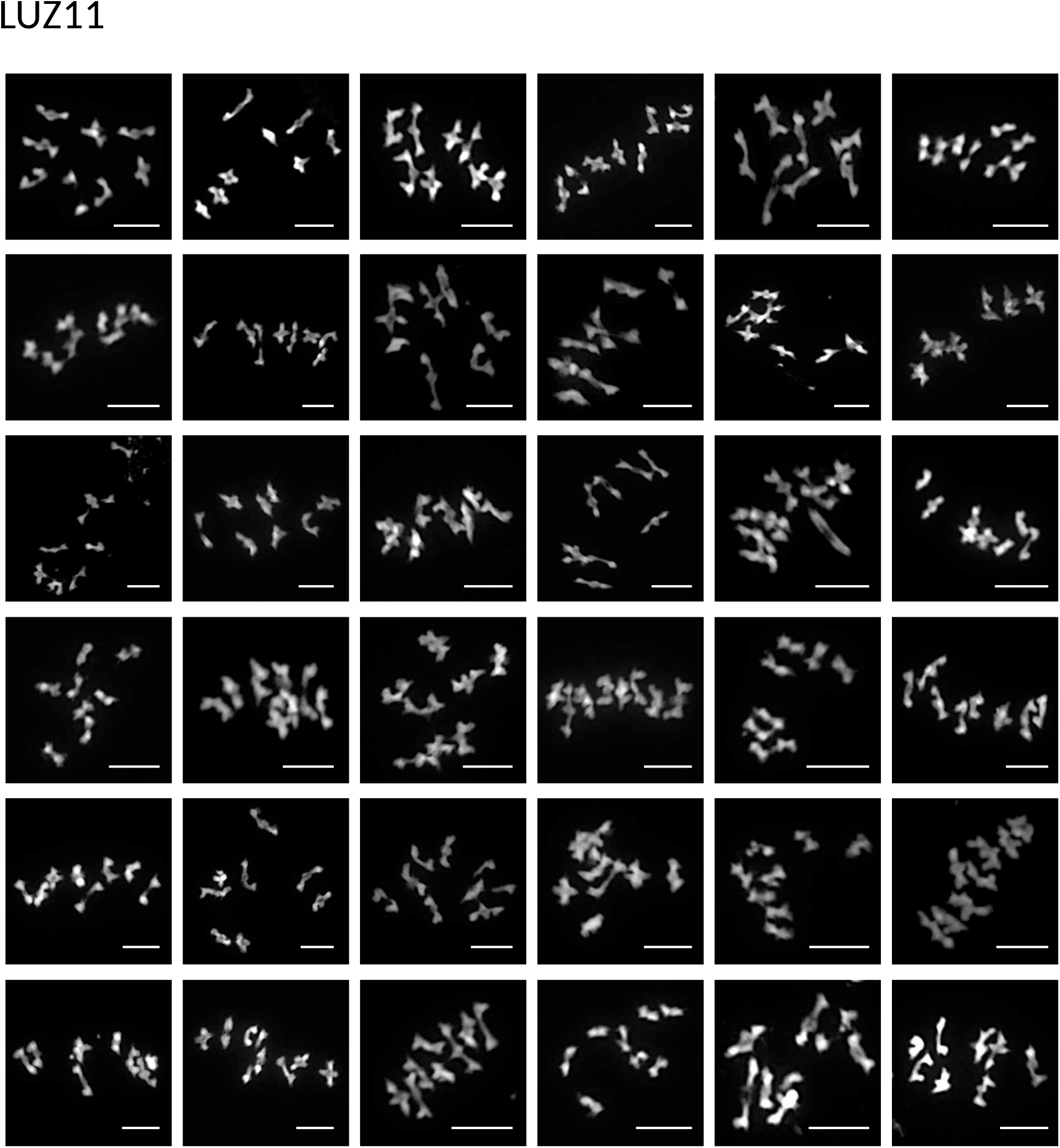
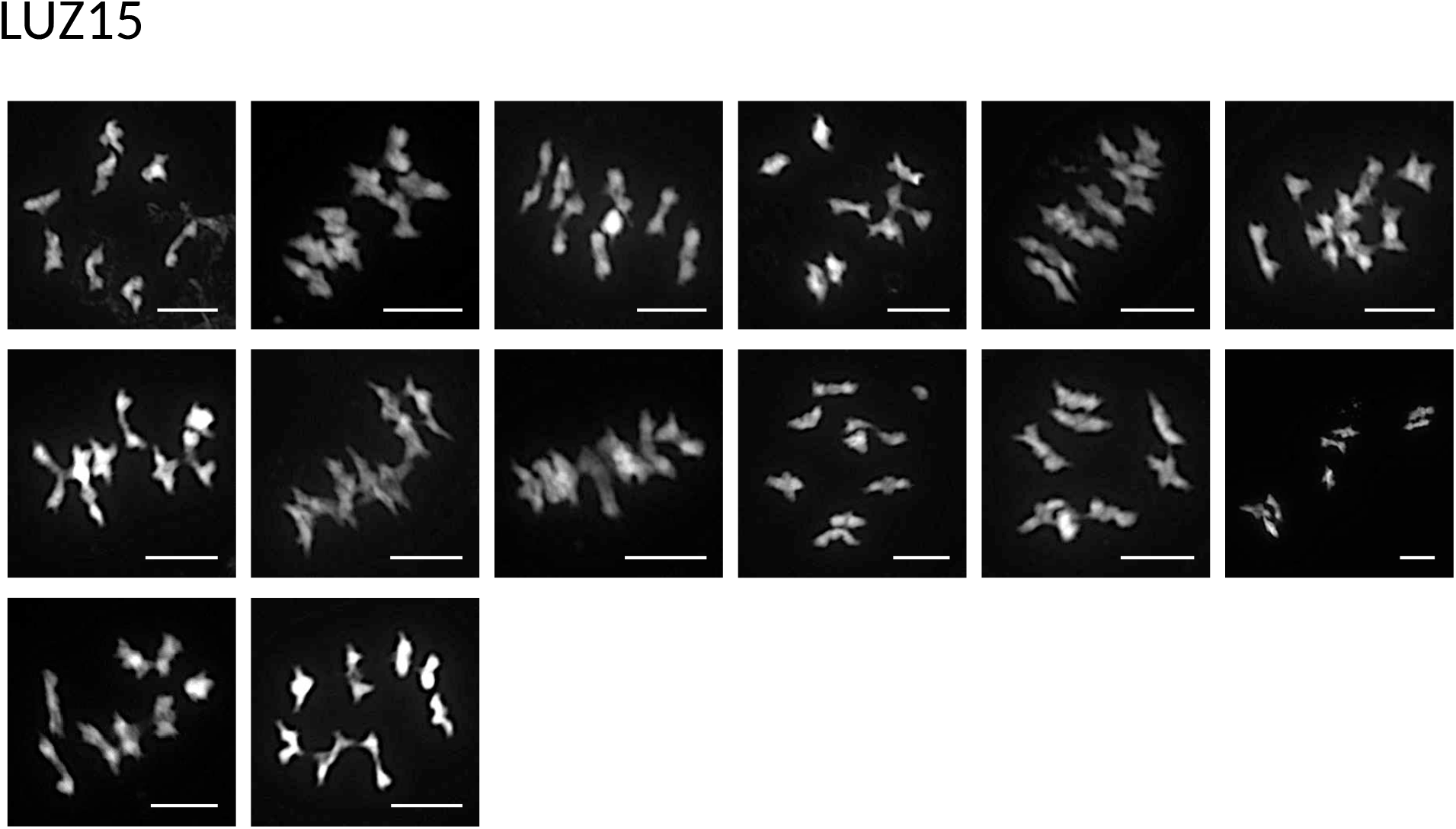
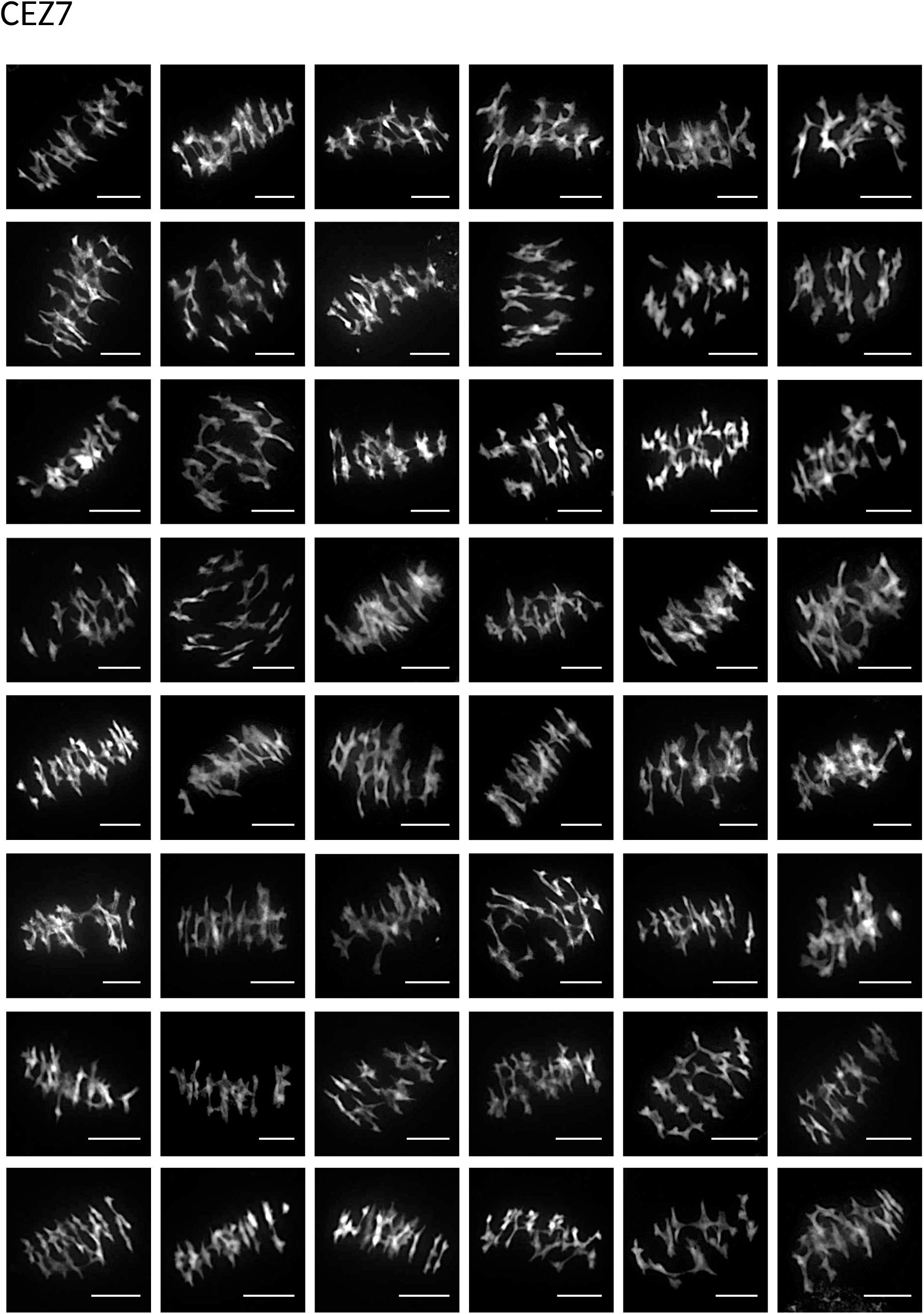
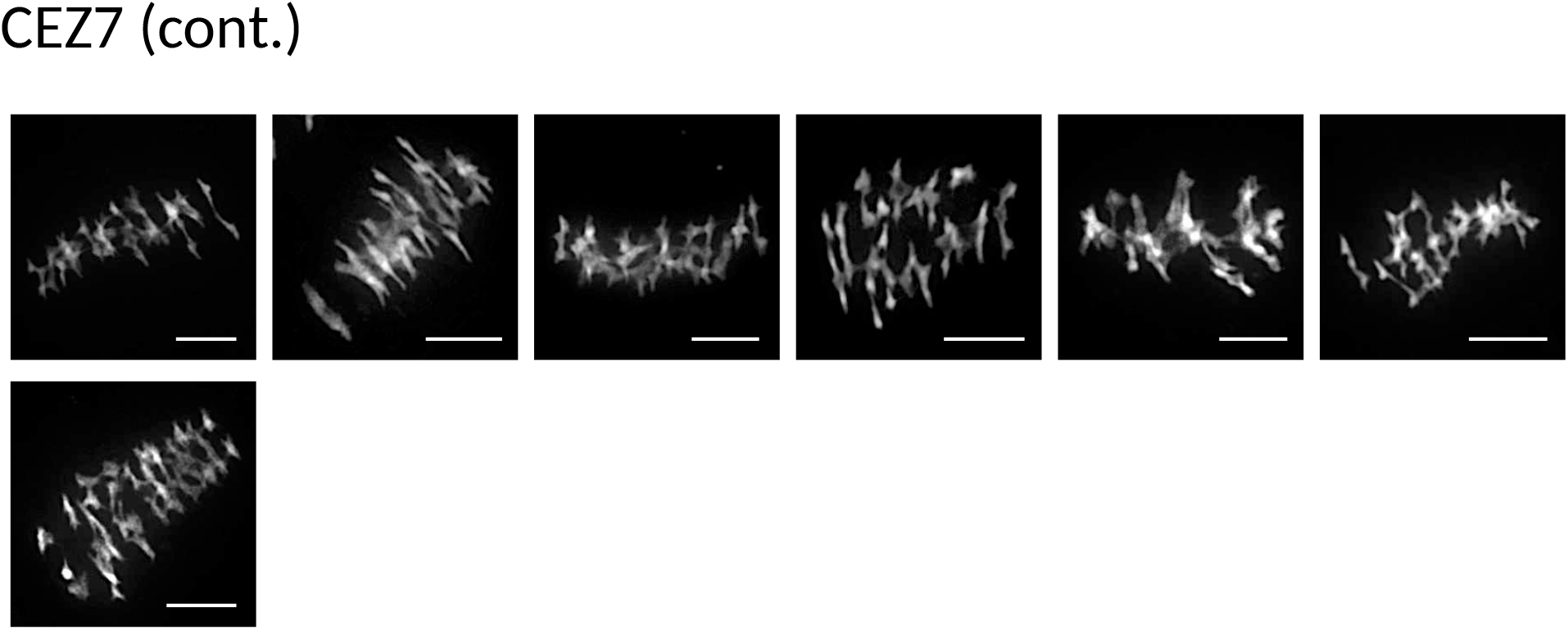
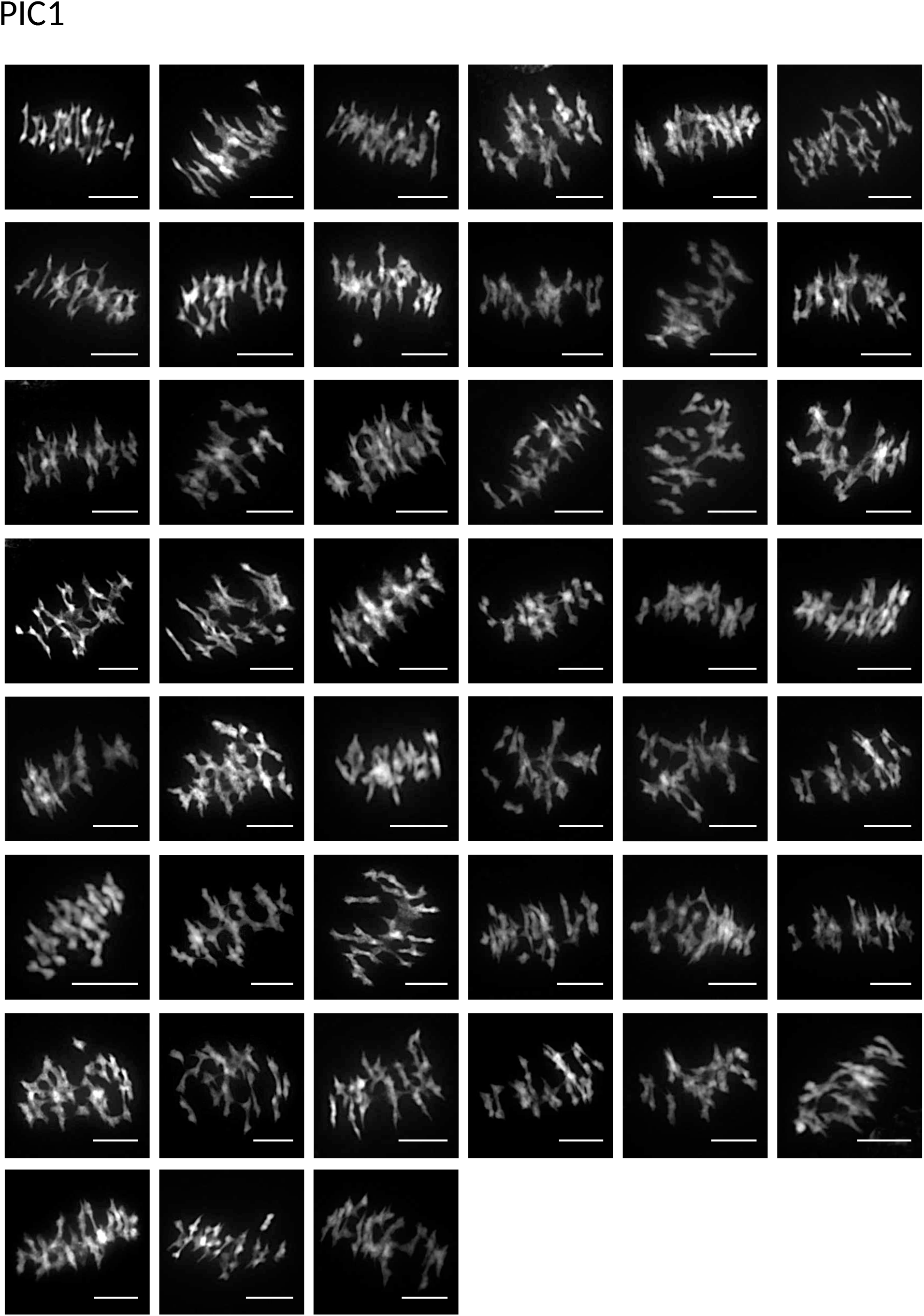
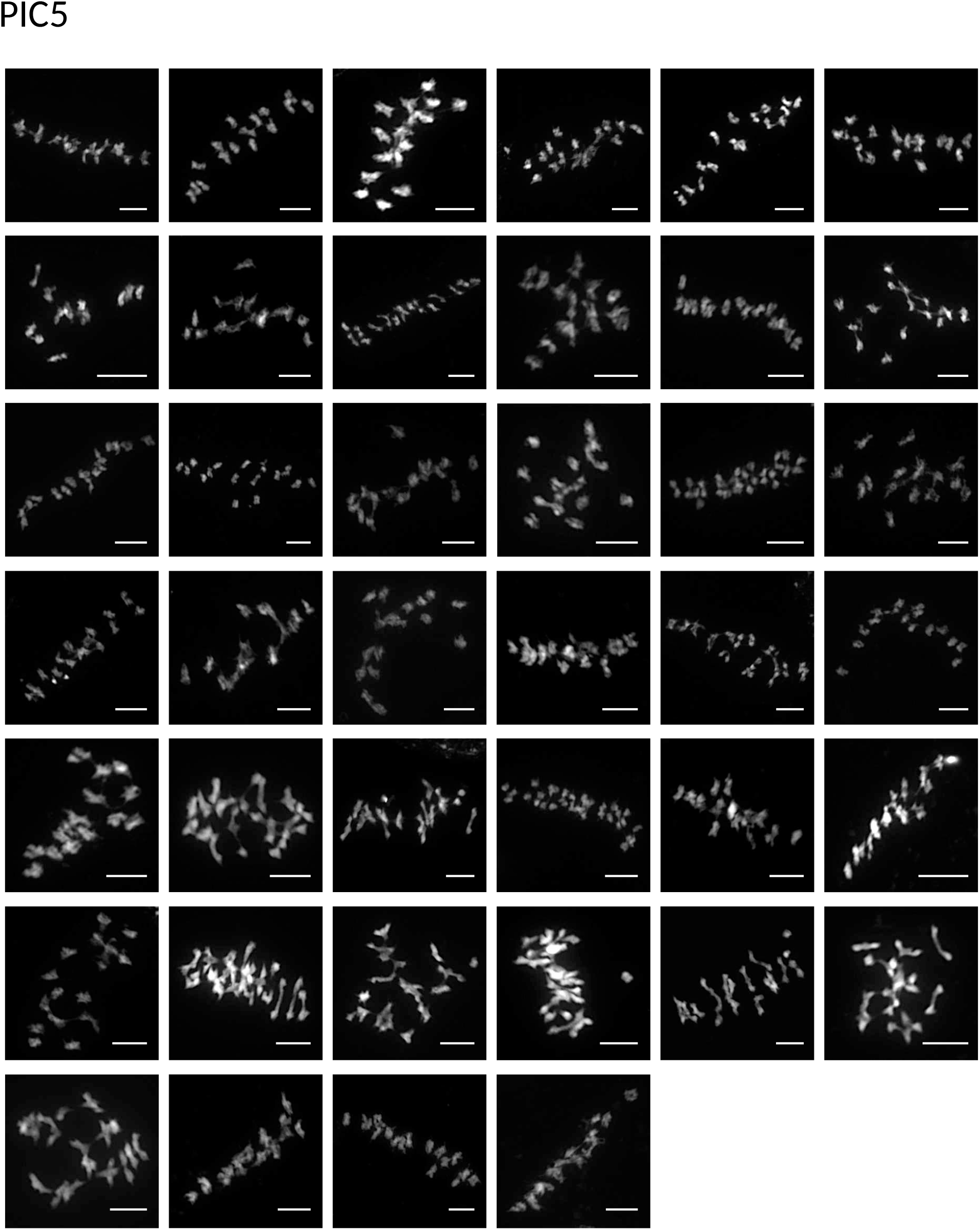
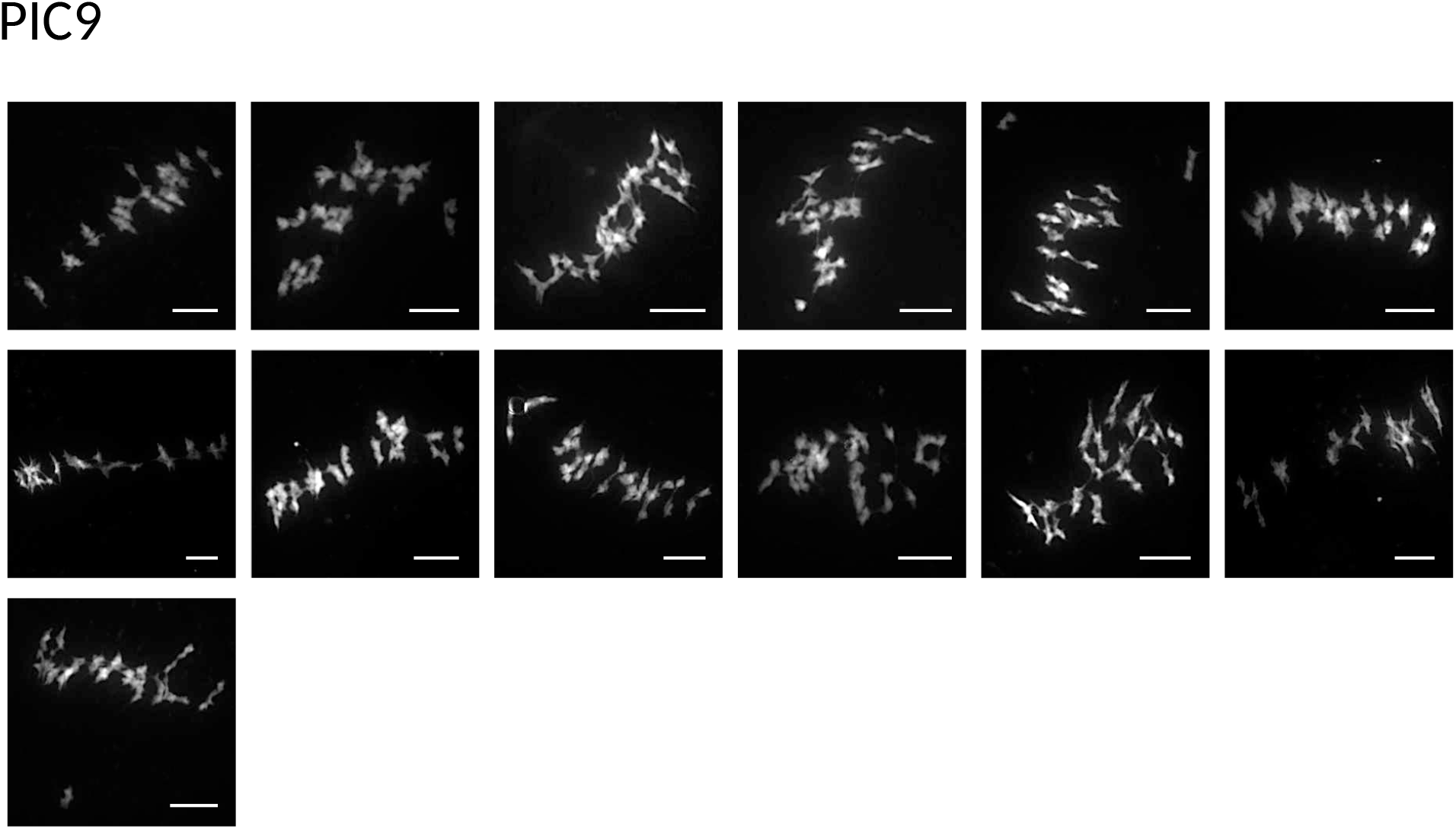
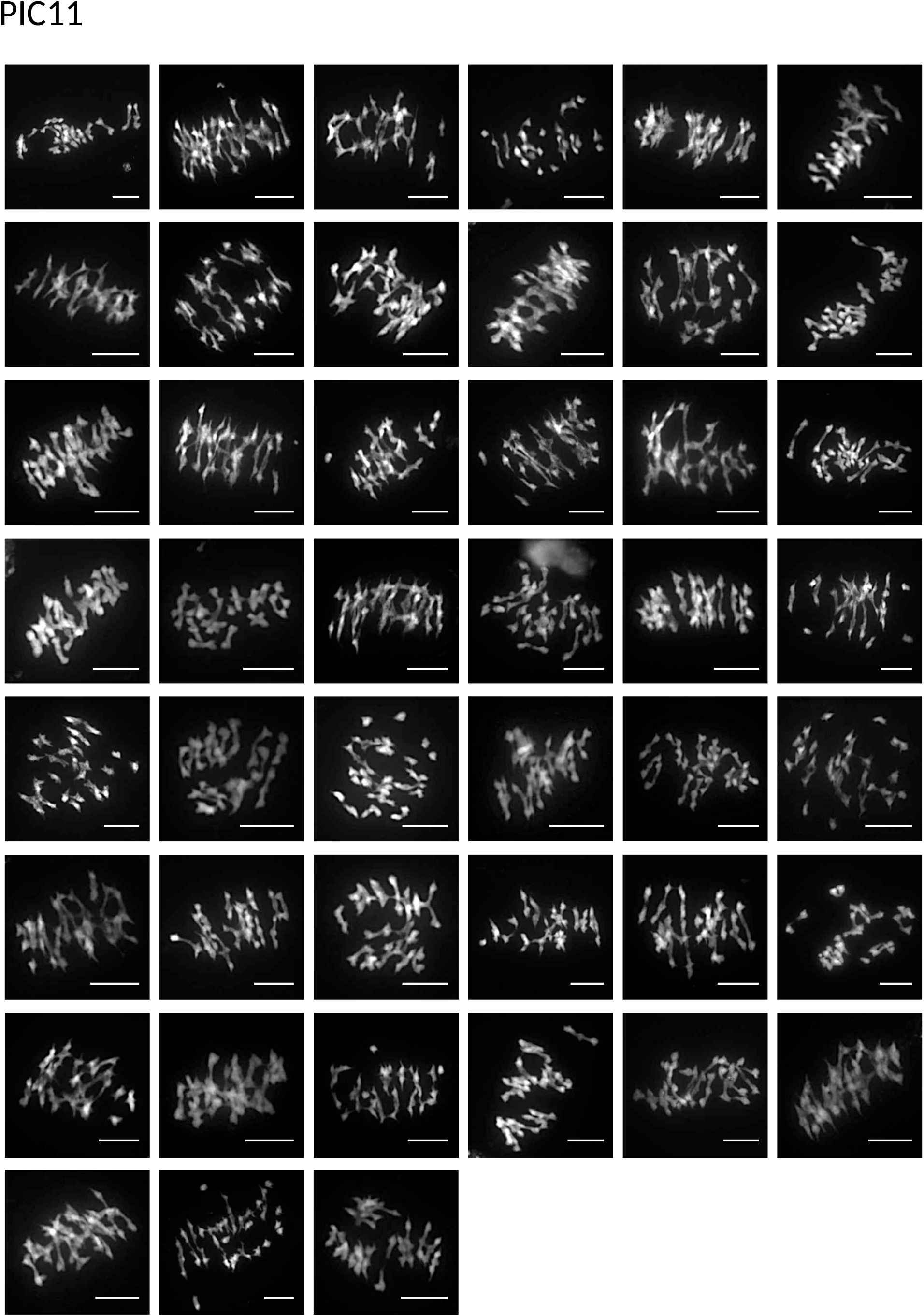
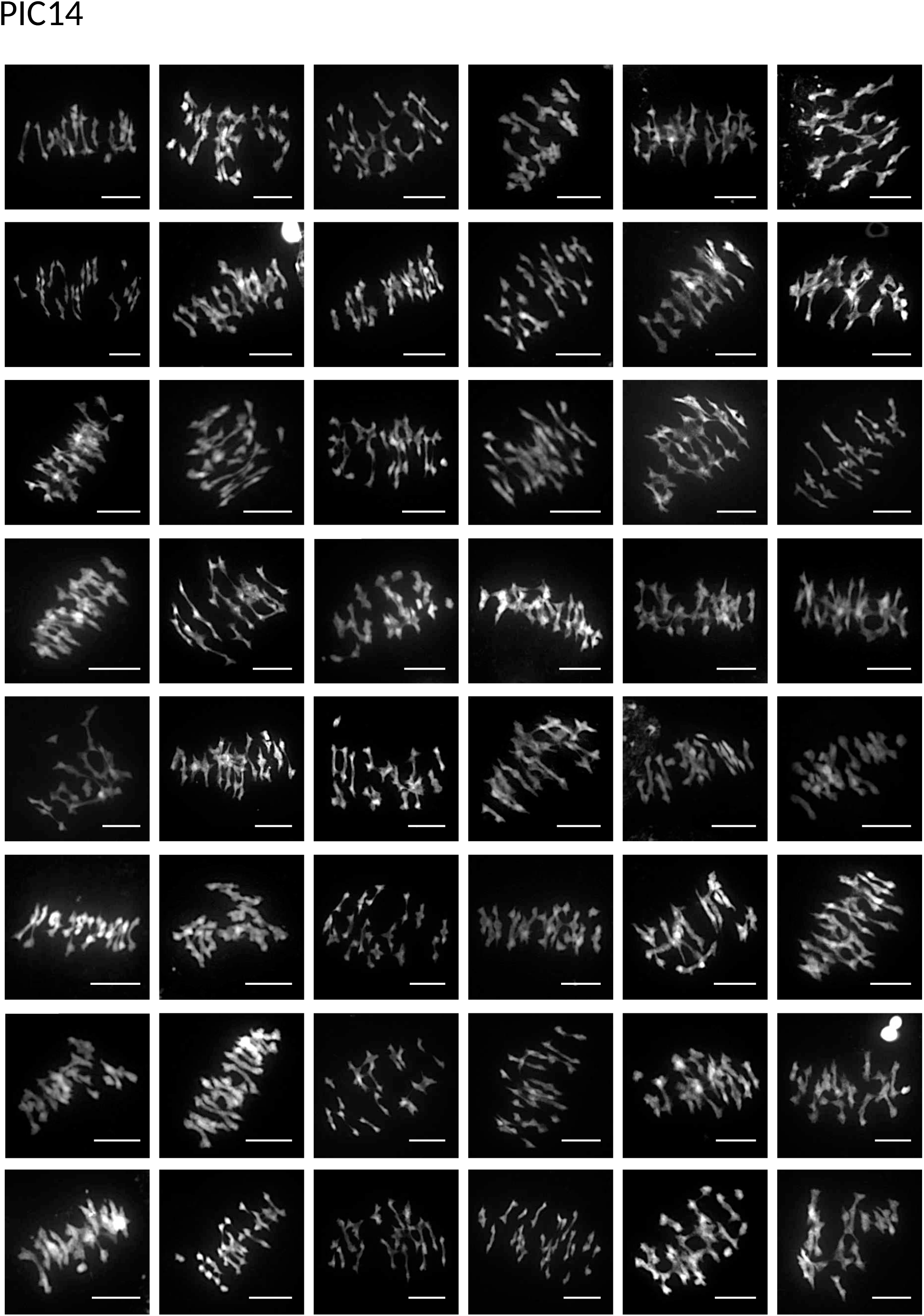
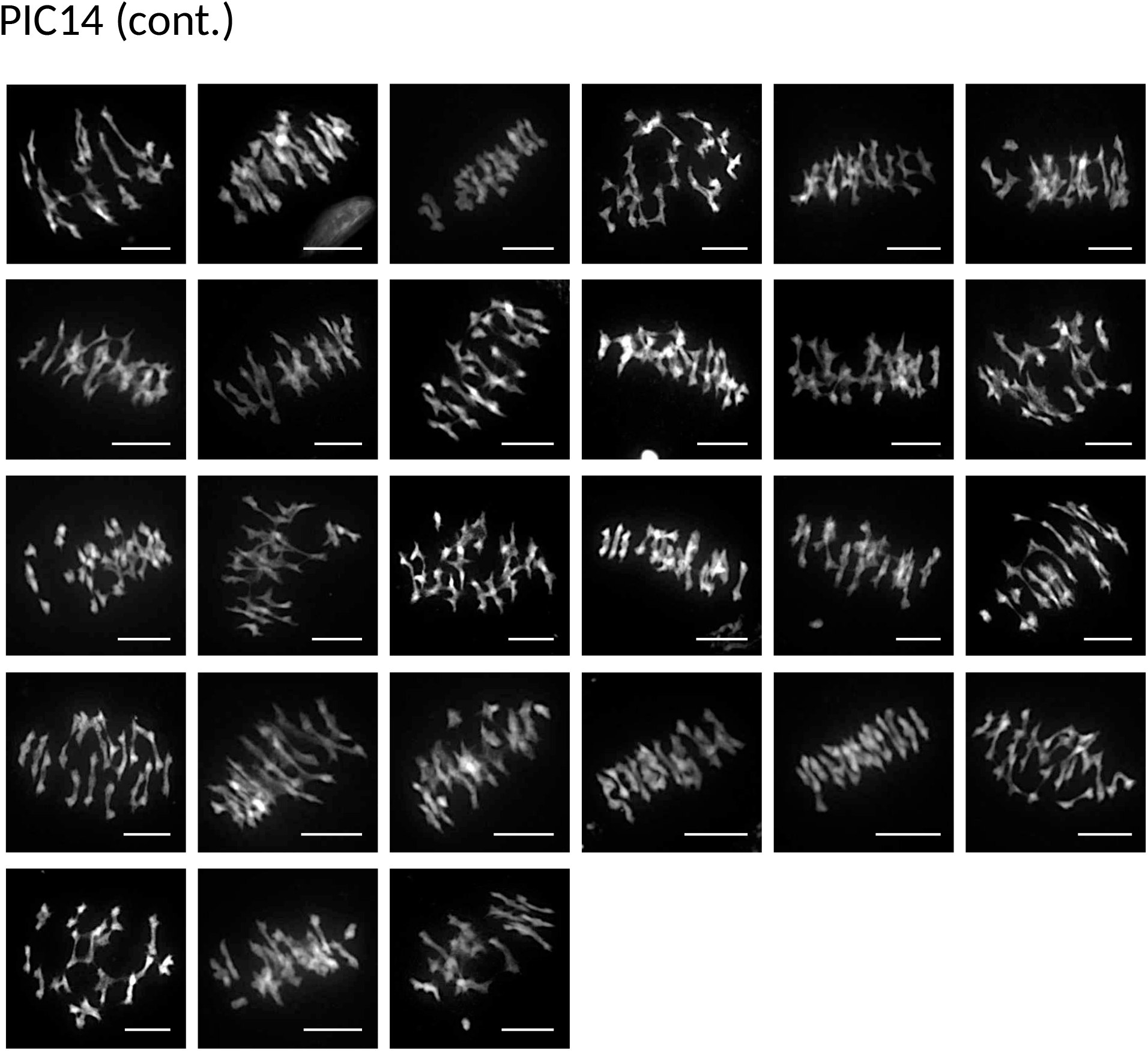
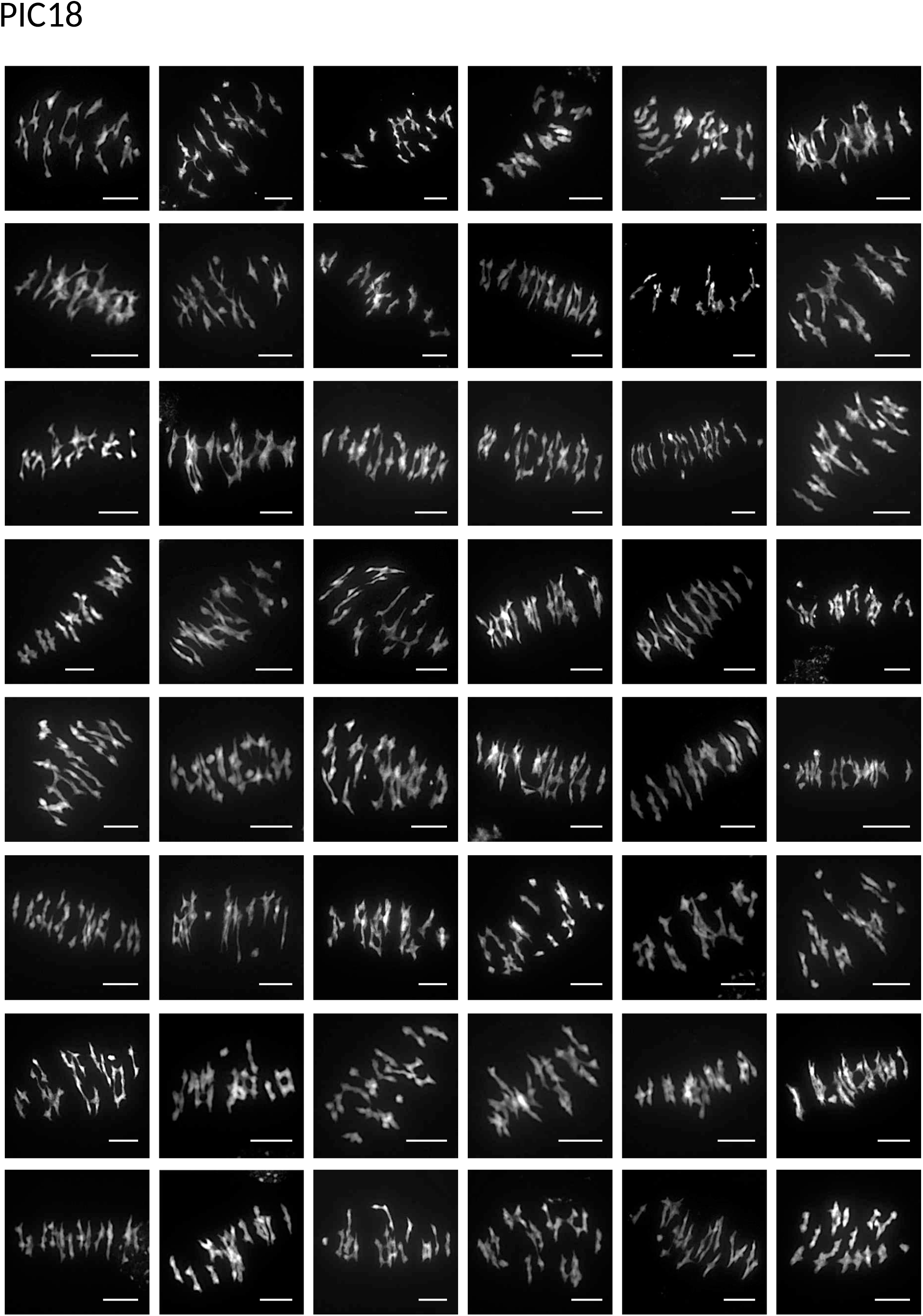
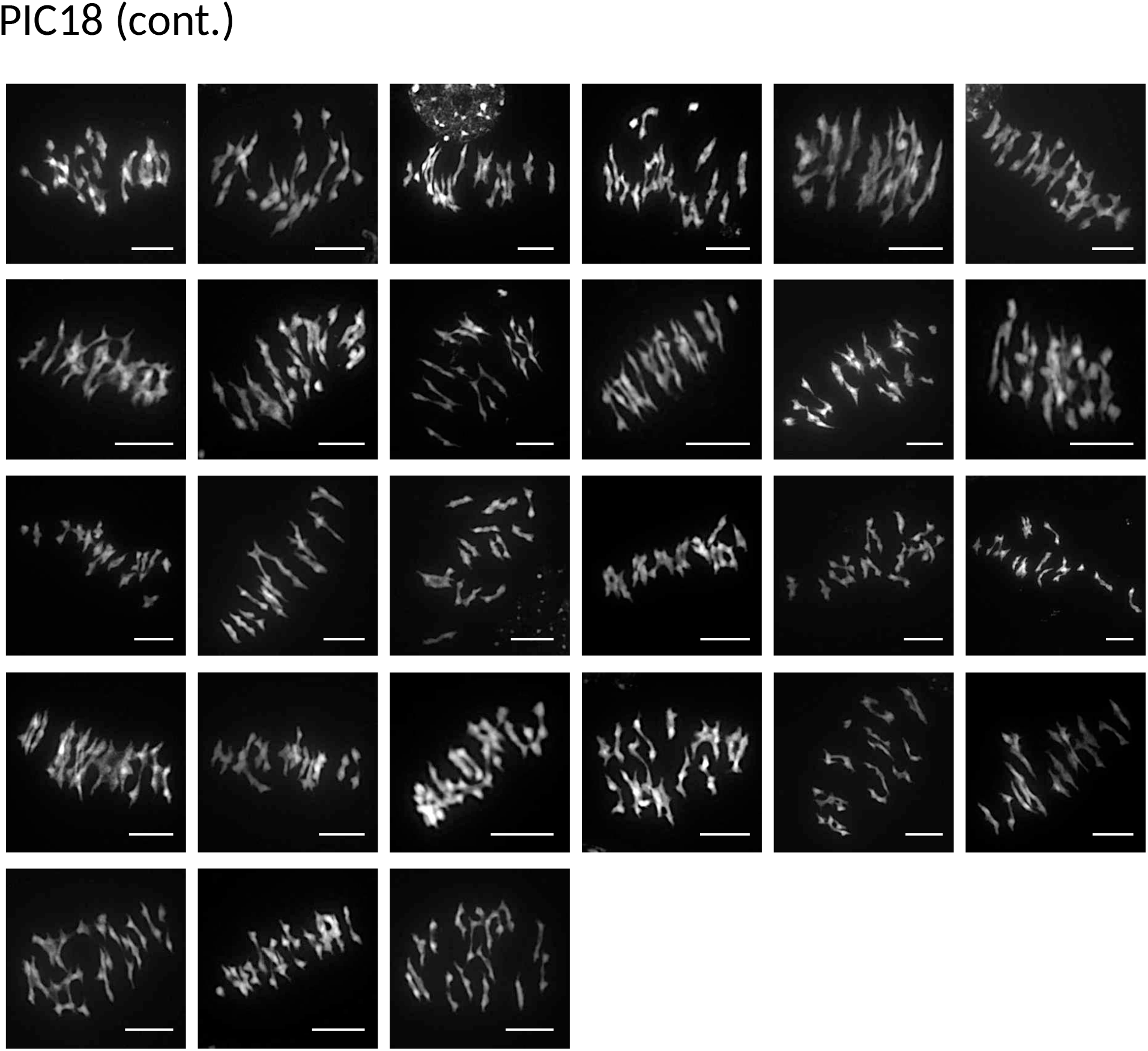
DAPI-stained meiotic (metaphase I) chromosomes of diploid (2n = 16; VKR6, VKR8, LUZ3, LUZ8, LUZ10, LUZ11, LUZ15) and tetraploid (2n = 32; CEZ7, PIC1, PIC5, PIC9, PIC11, PIC14, PIC1*8)* individuals of *Cardamine amara*. Scale bars = 10 μm.

## Other supplementary material

### Additional data Tables (separate files)

**Table S1**. GPS coordinates of population localities.

**Table S2**. Mean depth of coverage (MDOC) per pool of individuals from each population.

**Table S3**. Genes in the top 1% of Fst scores (1000 bp windows) in C. amara. Note: red lines denote six genes which are candidates also in A. arenosa.

**Table S4**. GO terms enriched in C. amara WGD candidate genes. Annotated: # genes in the GO category, Significant: # candidate genes in each category, p-values crom Fisher’s exact test (’elim’ method).

**Table S5**. Top 1% of amino acid substitutions with the highest fineMAV score.

**Table S6**. Genes in the top 1% of Fst scores (1000 bp windows) in A. arenosa.

**Table S7**. GO terms enriched in A. arenosa WGD candidate genes. Annotated: # genes in the GO category, Significant: # candidate genes in each category, p-values crom Fisher’s exact test (’elim’ method).

**Table S8.** Targeted search for patterns suggesting directional selection in C. amara orthologs of candidate A. arenosa meiosis genes.

**Table S9.** Chromosome stability scoring of individual diploid (2n = 16) and autotetraploid (2n = 32) plants of *Cardamine amara* at meiotic metaphase I.

**Table S10.** *C. amara* candidate genes that have more than one associated protein among *A. arenosa* candidates.

**Table S11.** Quality checks of DNA isolated from LUZ.

**Table S12.** Assessment of genome completeness using BUSCO.

## References

1. Y. Van De Peer, E. Mizrachi, K. Marchal, The evolutionary significance of polyploidy. Nat. Rev. Genet. 18, 411–424 (2017).

2. K. Bomblies, A. Madlung, Polyploidy in the Arabidopsis genus. Chromosom. Res. 22, 117–134 (2014).

3. L. Yant, K. Bomblies, Genome management and mismanagement—cell-level opportunities and challenges of whole-genome duplication. Genes Dev. 29, 2405–2419 (2015).

4. A. Lloyd, K. Bomblies, Meiosis in autopolyploid and allopolyploid Arabidopsis. Curr. Opin. Plant Biol. 30, 116–122 (2016).

5. D. Y. Chao, et al., Polyploids exhibit higher potassium uptake and salinity tolerance in Arabidopsis. Science (80-.). 341, 658–659 (2013).

6. J. J. Doyle, J. E. Coate, Polyploidy, the nucleotype, and novelty: The impact of genome doubling on the biology of the cell. Int. J. Plant Sci. 180, 1–52 (2019).

7. A. M. Selmecki, et al., Polyploidy can drive rapid adaptation in yeast. Nature 519, 349–351 (2015).

8. P. Monnahan, et al., Pervasive population genomic consequences of genome duplication in Arabidopsis arenosa. Nat. Ecol. Evol. 3, 457–468 (2019).

9. S. Marburger, et al., Interspecific introgression mediates adaptation to whole genome duplication. Nat. Commun. (2019) https:/doi.org/10.1038/s41467-019-13159-5.

10. P. Baduel, S. Bray, M. Vallejo-Marin, F. Kolář, L. Yant, The “Polyploid Hop”: Shifting challenges and opportunities over the evolutionary lifespan of genome duplications. Front. Ecol. Evol. 6 (2018).

11. K. Bomblies, J. D. Higgins, L. Yant, Meiosis evolves: Adaptation to external and internal environments. New Phytol. 208, 306–323 (2015).

12. F. Kolář, et al., Northern glacial refugia and altitudinal niche divergence shape genome-wide differentiation in the emerging plant model Arabidopsis arenosa. Mol. Ecol. (2016) https:/doi.org/10.1111/mec.13721.

13. J. D. Hollister, et al., Genetic Adaptation Associated with Genome-Doubling in Autotetraploid Arabidopsis arenosa. PLoS Genet. 8 (2012).

14. L. Yant, et al., Meiotic adaptation to genome duplication in Arabidopsis arenosa. Curr. Biol. 23, 2151–2156 (2013).

15. P. J. Seear, et al., A novel allele of ASY3 is associated with greater meiotic stability in autotetraploid Arabidopsis lyrata. PLoS Genet. in press (2020).

16. X. C. Huang, D. A. German, M. A. Koch, Temporal patterns of diversification in Brassicaceae demonstrate decoupling of rate shifts and mesopolyploidization events. Ann. Bot. 125, 29–47 (2020).

17. J. Lihová, J. F. Aguilar, K. Marhold, G. N. Feliner, Origin of the disjunct tetraploid Cardamine amporitana (Brassicaceae) assessed with nuclear and chloroplast DNA sequence data. Am. J. Bot. 91, 1231–1242 (2004).

18. J. Zozomová-Lihová, et al., Cytotype distribution patterns, ecological differentiation, and genetic structure in a diploid–tetraploid contact zone of Cardamine amara. Am. J. Bot. 102, 1380–1395 (2015).

19. K. Marhold, M. Huthmann, H. Hurka, Evolutionary history of the polyploid complex of Cardamine amara (Brassicaceae): isozyme evidence. Plant Syst. Evol. 233, 15–28 (2002).

20. K. Marhold, Taxonomic evaluation of the tetraploid populations of Cardamine amara (Brassicaceae) from the Eastern Alps and adjacent areas. Bot. Helv. 109, 67–84 (1999).

21. N. Vijay, et al., Evolution of heterogeneous genome differentiation across multiple contact zones in a crow species complex. Nat. Commun. 7 (2016).

22. M. Szpak, et al., FineMAV: Prioritizing candidate genetic variants driving local adaptations in human populations. Genome Biol. 19 (2018).

23. K. M. Culligan, J. B. Hays, Arabidopsis MutS homologs - AtMSH2, AtMSH3, AtMSH6, and a novel AtMSH7 - Form three distinct protein heterodimers with different specificities for mismatched DNA. Plant Cell 12, 991–1002 (2000).

24. S. Y. Wu, K. Culligan, M. Lamers, J. Hays, Dissimilar mispair-recognition spectra of Arabidopsis DNA-mismatch-repair proteins MSH2·MSH6 (MutSα) and MSH2·MSH7 (MutSγ). Nucleic Acids Res. 31, 6027–6034 (2003).

25. X. Lu, et al., The Arabidopsis MutS homolog AtMSH5 is required for normal meiosis. Cell Res. 18, 589–599 (2008).

26. S. Panizza, T. Tanaka, A. Hochwagen, F. Eisenhaber, K. Nasmyth, Pds5 cooperates with cohesion in maintaining sister chromatid cohesion. Curr. Biol. 10, 1557–1564 (2000).

27. N. U. Siddiqui, P. E. Stronghill, R. E. Dengler, C. A. Hasenkampf, C. D. Riggs, Mutations in Arabidopsis condensin genes disrupt embryogenesis, meristem organization and segregation of homologous chromosomes during meiosis. Development 130, 3283–3295 (2003).

28. C. M. Liu, D. W. Meinke, The titan mutants of Arabidopsis are disrupted in mitosis and cell cycle control during seed development. Plant J. 16, 21–31 (1998).

29. H. Shaked, N. Avivi-Ragolsky, A. A. Levy, Involvement of the arabidopsis SWI2/SNF2 chromatin remodeling gene family in DNA damage response and recombination. Genetics 173, 985–994 (2006).

30. J. Ogas, S. Kaufmann, J. Henderson, C. Somerville, PICKLE is a CHD3 chromatin-remodeling factor that regulates the transition from embryonic to vegetative development in Arabidopsis. Proc. Natl. Acad. Sci. U. S. A. 96, 13839–13844 (1999).

31. E. Perruc, N. Kinoshita, L. Lopez-Molina, The role of chromatin-remodeling factor PKL in balancing osmotic stress responses during Arabidopsis seed germination. Plant J. 52, 927–936 (2007).

32. X. Kang, et al., HRB2 and BBX21 interaction modulates Arabidopsis ABI5 locus and stomatal aperture. Plant Cell Environ. 41, 1912–1925 (2018).

33. E. Aichinger, C. B. R. Villar, R. di Mambro, S. Sabatini, C. Köhler, The CHD3 chromatin remodeler PICKLE and polycomb group proteins antagonistically regulate meristem activity in the Arabidopsis root. Plant Cell 23, 1047–1060 (2011).

34. Y. Jing, Q. Guo, R. Lin, The Chromatin-Remodeling Factor PICKLE Antagonizes Polycomb Repression of FT to Promote Flowering. Plant Physiol. 181, 656–668 (2019).

35. P. Bundock, P. Hooykaas, An Arabidopsis hAT-like transposase is essential for plant development. Nature 436, 282–284 (2005).

36. K. Tamura, Y. Adachi, K. Chiba, K. Oguchi, H. Takahashi, Identification of Ku70 and Ku80 homologues in Arabidopsis thaliana: Evidence for a role in the repair of DNA double-strand breaks. Plant J. 29, 771–781 (2002).

37. M. Leiden U. Knip, “Daysleeper : from genomic parasite to indispensable gene.” (2012).

38. J. Glover, M. Grelon, S. Craig, A. Chaudhury, E. Dennis, Cloning and characterization of MS5 from Arabidopsis: A gene critical in male meiosis. Plant J. 15, 345–356 (1998).

39. M. Cifuentes, et al., TDM1 Regulation Determines the Number of Meiotic Divisions. PLoS Genet. 12 (2016).

40. D. Kawa, et al., SnRK2 Protein Kinases and mRNA Decapping Machinery Control Root Development and Response to Salt. Plant Physiol. 182, 361–377 (2020).

41. H. Fujii, P. E. Verslues, J. K. Zhu, Arabidopsis decuple mutant reveals the importance of SnRK2 kinases in osmotic stress responses in vivo. Proc. Natl. Acad. Sci. U. S. A. 108, 1717–1722 (2011).

42. S. R. Cutler, P. L. Rodriguez, R. R. Finkelstein, S. R. Abrams, Abscisic Acid: Emergence of a Core Signaling Network. Annu. Rev. Plant Biol. 61, 651–679 (2010).

43. M. E. Williams, et al., Mutations in the Arabidopsis phosphoinositide phosphatase gene SAC9 lead to overaccumulation of PtdIns(4,5)P2 and constitutive expression of the stress-response pathway. Plant Physiol. 138, 686–700 (2005).

44. B. C. Meyers, M. Morgante, R. W. Michelmore, TIR-X and TIR-NBS proteins: Two new families related to disease resistance TIR-NBS-LRR proteins encoded in Arabidopsis and other plant genomes. Plant J. 32, 77–92 (2002).

45. A. V. Klepikova, A. S. Kasianov, E. S. Gerasimov, M. D. Logacheva, A. A. Penin, A high resolution map of the Arabidopsis thaliana developmental transcriptome based on RNA-seq profiling. Plant J. 88, 1058–1070 (2016).

46. H. H. Kunz, et al., Plastidial transporters KEA1, −2, and −3 are essential for chloroplast osmoregulation, integrity, and pH regulation in Arabidopsis. Proc. Natl. Acad. Sci. U. S. A. 111, 7480–7485 (2014).

47. A. B. Stephan, H. H. Kunz, E. Yang, J. I. Schroeder, Rapid hyperosmotic-induced Ca2+ responses in Arabidopsis thaliana exhibit sensory potentiation and involvement of plastidial KEA transporters. Proc. Natl. Acad. Sci. U. S. A. 113, E5242–E5249 (2016).

48. J. M. Colmenero-Flores, et al., Identification and functional characterization of cation-chloride cotransporters in plants. Plant J. 50, 278–292 (2007).

49. Hejný Slavomil, Slavík Bohumil, Květena České republiky 3 (Academia, 2003).

50. D. Szklarczyk, et al., STRING v10: Protein-protein interaction networks, integrated over the tree of life. Nucleic Acids Res. 43, D447–D452 (2015).

51. R. R. Hudson, J. A. Coyne, Mathematical consequences of the genealogical species concept. Evolution (N. Y). 56, 1557–1565 (2002).

52. M. Seppey, M. Manni, E. M. Zdobnov, “BUSCO: Assessing genome assembly and annotation completeness” in Methods in Molecular Biology, (2019), pp. 227–245.

53. J. B. Spalding, P. J. Lammers, BLAST Filter and GraphicAlign: Rule-based formation and analysis of sets of related DNA and protein sequences. Nucleic Acids Res. 32 (2004).

54. G. Marçais, et al., MUMmer4: A fast and versatile genome alignment system. PLoS Comput. Biol. 14 (2018).

55. M. Stanke, S. Waack, Gene prediction with a hidden Markov model and a new intron submodel in Bioinformatics, (2003).

56. E. Quevillon, et al., InterProScan: Protein domains identifier. Nucleic Acids Res. 33 (2005).

57. A. Conesa, S. Götz, Blast2GO: A comprehensive suite for functional analysis in plant genomics. Int. J. Plant Genomics 2008 (2008).

58. E. D.M., K. S., OrthoFinder2: fast and accurate phylogenomic orthology analysis from gene sequences. bioRxiv, 466201 (2018).

59. M. Nei, Genetic Distance between Populations. Am. Nat. (1972) https:/doi.org/10.1086/282771.

60. S. Anand, et al., Next generation sequencing of pooled samples: Guideline for variants’ filtering. Sci. Rep. 6 (2016).

61. T. Jombart, I. Ahmed, adegenet 1.3-1: New tools for the analysis of genome-wide SNP data. Bioinformatics 27, 3070–3071 (2011).

62. P. Rentzsch, D. Witten, G. M. Cooper, J. Shendure, M. Kircher, CADD: Predicting the deleteriousness of variants throughout the human genome. Nucleic Acids Res. 47, D886–D894 (2019).

63. R. Grantham, Amino acid difference formula to help explain protein evolution. Science (80-.). 185, 862–864 (1974).

64. A. McKenna, et al., The genome analysis toolkit: A MapReduce framework for analyzing next-generation DNA sequencing data. Genome Res. 20, 1297–1303 (2010).

65. A. Alexa, J. Rahnenführer, Gene set enrichment analysis with topGO. Bioconductor Improv. (2007).

66. D. Smedley, et al., BioMart - Biological queries made easy. BMC Genomics (2009) https:/doi.org/10.1186/1471-2164-10-22.

67. S. Grossmann, S. Bauer, P. N. Robinson, M. Vingron, Improved detection of overrepresentation of Gene-Ontology annotations with parent-child analysis. Bioinformatics 23, 3024–3031 (2007).

68. R. R Development Core Team, R: A Language and Environment for Statistical Computing (2011).

69. T. Mandáková, K. Marhold, M. A. Lysak, The widespread crucifer species Cardamine flexuosa is an allotetraploid with a conserved subgenomic structure. New Phytol. 201, 982–992 (2014).

## Supplemental References

1. M. Martin, Cutadapt removes adapter sequences from high-throughput sequencing reads. EMBnet.journal 17, 10(2011).

2. N. Joshi, J. Fass, Sickle: A sliding-window, adaptive, quality-based trimming tool for FastQ files (Version 1.33) [Software]. Available at https://github.com/najoshi/sickle., 2011 (2011).

3. H. Li, et al., The Sequence Alignment/Map format and SAMtools. Bioinformatics 25, 2078–2079 (2009).

4. H. Li, R. Durbin, Fast and accurate short read alignment with Burrows-Wheeler transform. Bioinformatics 25, 1754–1760 (2009).

5. Broad Institute, Picard Tools - By Broad Institute. Github (2009).

6. A. McKenna, et al., The genome analysis toolkit: A MapReduce framework for analyzing next-generation DNA sequencing data. Genome Res. 20, 1297–1303 (2010).

7. E. Garrison, G. Marth, Haplotype-based variant detection from short-read sequencing (2012).

8. V. Narasimhan, et al., BCFtools/RoH: A hidden Markov model approach for detecting autozygosity from next-generation sequencing data. Bioinformatics (2016) https:/doi.org/10.1093/bioinformatics/btw044.

9. L. Venturini, S. Caim, G. G. Kaithakottil, D. L. Mapleson, D. Swarbreck, Leveraging multiple transcriptome assembly methods for improved gene structure annotation. Gigascience 7 (2018).

10. P. Cingolani, et al., A program for annotating and predicting the effects of single nucleotide polymorphisms, SnpEff: SNPs in the genome of Drosophila melanogaster strain w1118; iso-2; iso-3. Fly (Austin). 6, 80–92 (2012).

11. M. Szpak, et al., FineMAV: Prioritizing candidate genetic variants driving local adaptations in human populations. Genome Biol. 19 (2018).

